# The interferon-gamma response pathway is a targetable driver of FSHD muscular dystrophy progression

**DOI:** 10.64898/2026.10.13.758792

**Authors:** Beatrice Biferali, Federica Del Prete, Mara Salomè, Jose Manuel Garcia Manteiga, Reza Alemohammad, Miriam Mombellardo, Maria Pannese, Emanuele Mocciaro, Francesca Torri, İlker Eren, Elisabetta Ferraro, Dicle Ballikaya, Beatrice Ciurli, Arianna Votta, Daniele Campolungo, Daniel C. Turner, Monica Canepari, Roberto Bottinelli, Gabriele Siciliano, Serpil Eraslan, Giulia Ricci, Davide Gabellini

## Abstract

Facioscapulohumeral muscular dystrophy (FSHD) is a leading neuromuscular disorder caused by the aberrant expression of the transcription factor double homeobox 4 (DUX4) in skeletal muscle. While DUX4 is normally restricted to early embryonic development, its reactivation in FSHD triggers chronic inflammation and progressive muscle degeneration. However, the precise molecular links between DUX4 and persistent immune activation remain poorly understood. Here, we show that DUX4 induces a viral mimicry-like pathway that is fundamentally uncoupled from canonical type I interferon (IFN) activation. Instead, DUX4 establishes a non-canonical innate immune state dominated by IFN-γ signaling, providing a mechanistic explanation for the non-resolving inflammation characteristic of the disease. We demonstrate that while IFN-γ treatment alone is sufficient to recapitulate key DUX4 pathogenic effects, its inhibition rescues muscle inflammation and halts disease progression in FSHD models. These findings define a “skewed viral mimicry” process driven by a disease-specific transcription factor and identify IFN-γ as a primary therapeutic target for tissue-informed immunomodulation in muscular dystrophy.

**Teaser:** IFN-γ response is the main upregulated inflammatory pathway in FSHD muscle and represents a key driver of disease progression.

## INTRODUCTION

Facioscapulohumeral muscular dystrophy (FSHD), one of the most common neuromuscular diseases, is a chronic illness leading to significant lifetime morbidity. FSHD is caused by aberrant expression of the transcription factor double homeobox 4 (DUX4) (1). DUX4 expression is normally confined to early embryonic development, while it is subsequently maintained silent in most somatic tissues, including skeletal muscle. FSHD is characterized by loss of silencing and aberrant reactivation of DUX4, which is toxic to the muscle tissue [reviewed in (1)].

The molecular mechanism through which DUX4 leads to FSHD is poorly known and, despite several clinical trials, there continues to be no cure or therapeutic treatment available to FSHD patients [reviewed in (1,2)].

Aberrant DUX4 expression has been associated with the activation of a large and still increasing list of potentially pathogenic effects including inhibition of myogenic differentiation, muscle atrophy, altered RNA processing, protein aggregation, immune response, and cell death [reviewed in (1)]. However, the involvement of DUX4 in many of these processes has been documented only using artificial systems ectopically expressing DUX4 and validation in FSHD samples is still lacking (1).

Since it is unclear which of the DUX4 downstream pathways contributes to the disease or whether targeting them offers any therapeutic benefit, most of the current therapeutic activities are focused on developing ways to block DUX4 expression. Nevertheless, increasing evidence suggests that DUX4 activated pathways might become independent from DUX4 and self-sustaining (3–11). Therefore, the identification of targetable DUX4-activated disease mediators bears strong therapeutic relevance.

While DUX4 is central to FSHD, immune activation and inflammation are increasingly recognized as key drivers of disease progression. Inflammation is a prominent feature that takes place early on in FSHD and anticipates muscle atrophy and its fibro-fatty substitution (12), unlike other muscular dystrophies in which inflammation is generally secondary to muscle degeneration [reviewed in (13)]. FSHD muscles with signs of inflammation have a faster progression to fat replacement, and the rate of disease progression correlates with the severity of inflammation (12,14–16).

Despite the recognition that muscle inflammation is a prominent feature of FSHD, the underlying molecular mechanism remains poorly understood. Accordingly, the generic anti-inflammatory drugs tested so far have shown inconclusive effect on FSHD patients (17–22). The current research has largely overlooked the specific inflammatory processes in FSHD, particularly how inflammation may directly contribute to disease progression rather than serving merely as a byproduct of muscle degeneration. This is a critical gap, as understanding the inflammatory mechanism in FSHD could lead to the development of targeted therapeutic interventions.

Recent data suggest that an immune response to “FSHD antigens” might have a role in disease progression (23). However, no disease-specific autoantibodies have been detected in FSHD (24). Notably, FSHD muscle cells induce an innate immune response and immune-dependent FSHD muscle loss (25), further supporting the idea that inflammation directly contributes to muscle wasting in FSHD. Nevertheless, the nature of the “FSHD antigens” is currently unknown.

Here, we show that DUX4 induces an interferon gamma (IFN-γ) dependent inflammatory pathway playing a key role in muscle degeneration in FSHD. Importantly, we found that IFN-γ is sufficient to recapitulate several DUX4 toxic effects and is required for disease progression downstream of DUX4. Thus, our results point to an original mechanism responsible for muscle inflammation in FSHD and open the possibility to the development of novel therapeutic approaches to mitigate muscle degeneration and improve FSHD patient outcomes.

## RESULTS

### The IFN-γ response pathway is the most upregulated inflammatory pathway in FSHD muscles

To begin defining the molecular mechanism underlying FSHD muscle inflammation, we analyzed publicly available datasets comparing the expression profile of skeletal muscles biopsies from FSHD patients and healthy donors, or from the ACTA1-MCM;FLExDUX4 mouse model of FSHD (called MCM;FLExD from here onward) and ACTA1-MCM control mice (23,26–31). We found that the IFN-γ response is the most significantly upregulated inflammatory pathway in muscles of FSHD patients or MCM;FLExD mice compared to their controls (Fig. 1A) (Tables S1-3). Of note, the IFN-γ response pathway was also significantly upregulated in inflamed compared to not inflamed FSHD muscles, even within the same FSHD patient (Fig. 1A) (Tables S2-3).

**Fig. 1.**
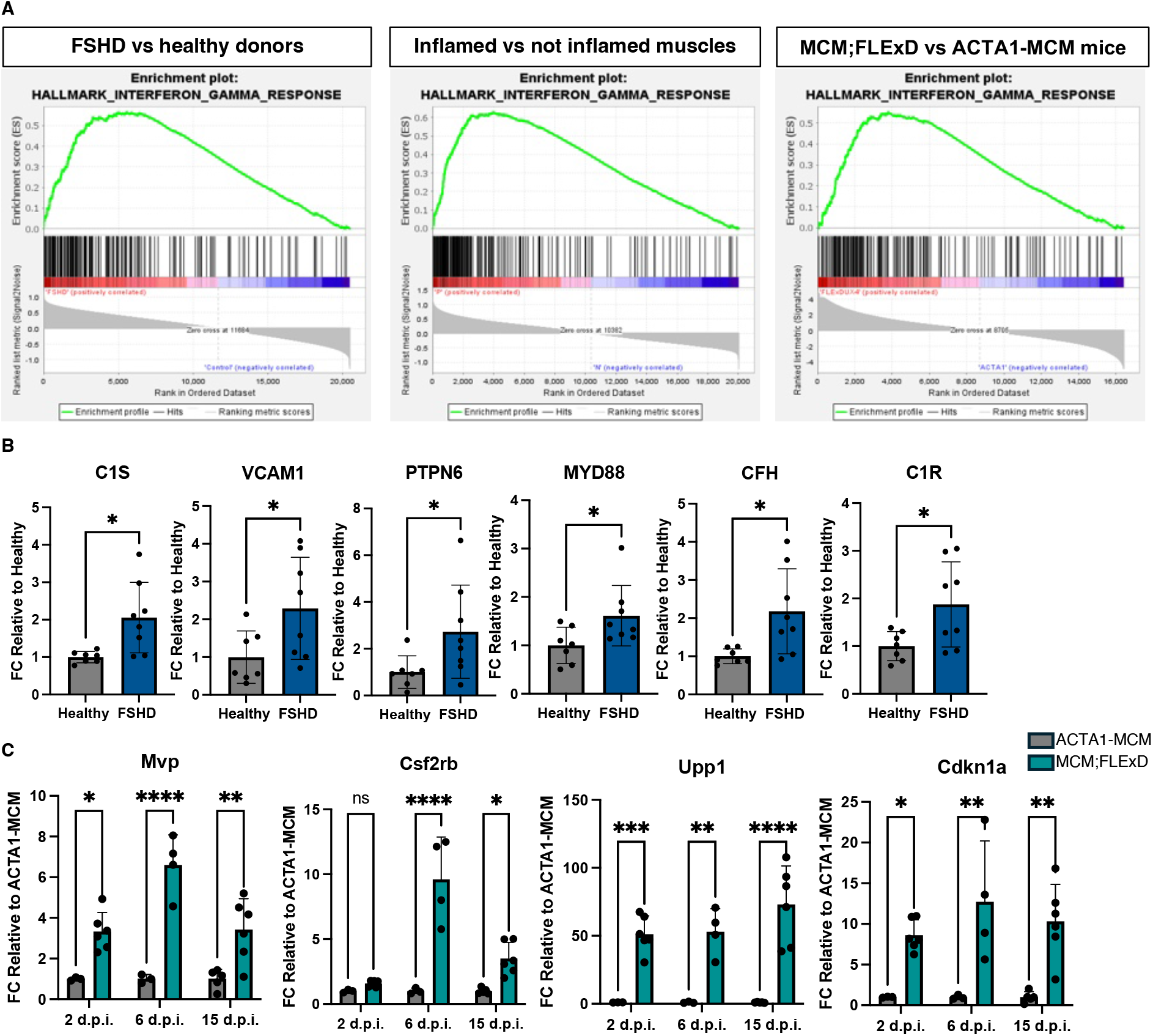
Activation of the IFN-γ response pathway in FSHD muscles. **(A)** Representative GSEA results showing activation of the IFN-γ response pathway in muscles of FSHD patients compared to healthy donors (left), inflamed FSHD muscles compared to not inflamed FSHD muscles (center), and muscles from MCM;FLExD compared to ACTA1-MCM mice (right). **(B)** RT-qPCR showing activation of the indicated IFN-γ response pathway genes in differentiated human primary muscle cells from FSHD patients (n=8) compared to healthy donors (n=7). Values expressed as fold change relative to healthy donor cells. Statistical significance assessed by Unpaired t-test, *p<0.05. **(C)** RT-qPCR showing activation of the indicated IFN-γ response pathway genes in muscle biopsies of MCM;FLExD mice compared to ACTA1-MCM mice at 2-6-15 d.p.i. n=4-6. Values expressed as fold change relative to ACTA1-MCM mice for each time point. Statistical significance assessed by two-way ANOVA, with Sidak’s multiple comparisons test, *p<0.05; **p<0.01; ***p<0.001, ****p<0.0001.

Intriguingly, we found a significant upregulation of genes specifically belonging to the IFN-γ response pathway in terminally differentiated primary muscle cells from FSHD patients compared to healthy donors (Table S4) (Fig. 1B). Beside validating our finding, this result indicates that activation of the IFN-γ response pathway is intrinsic to FSHD muscle cells.

MCM;FLExD is the most widely used FSHD animal model in academia and by companies developing FSHD therapeutics (30,32–43). Thanks to the skeletal muscle fiber-specific and tamoxifen (TMX) inducible ACTA1-MCM driver, in this model DUX4 expression is conditional, titratable, and restricted to myofibers (30). To induce DUX4, we used a low TMX regimen, which led to a very reproducible mild myopathy over the course of 15 days (Fig. S1). A mild disease phenotype allows us to maximize the possibility to model direct, primary effects of aberrant DUX4 expression. Notably, representative IFN-γ response pathway genes were significantly upregulated in muscles of MCM;FLExD compared to ACTA1-MCM mice already two days post DUX4 induction (d.p.i.) (Fig. 1C), when the level of muscle degeneration is extremely low (Fig. S1B) and (30).

### DUX4 is required and sufficient to activate a viral mimicry-like pathway

The majority of direct DUX4 targets are endogenous retroviruses and other repetitive elements (RE) (44,45). RE activation by DUX4 leads to the production of double stranded RNA (dsRNA), which accumulates in FSHD muscle cells and significantly contributes to DUX4 toxicity (44,45). In the cancer setting, dsRNA accumulation following RE activation is detected by pattern recognition dsRNA receptors (TLR-3, MDA5, RIG-I) and induces an innate immune “viral mimicry” response culminating with the activation of IFN signaling [reviewed in (46,47)]. Whether this pathway is activated or plays any role in skeletal muscle is currently unknown. Intriguingly, we found significant upregulation of dsRNA receptors, *IFN-γ* and other inflammatory molecules in terminally differentiated primary muscle cells of FSHD patients compared to healthy donors (Fig. 2A), supporting the activation of a viral mimicry-like pathway in FSHD muscle. Moreover, *DUX4* knockdown in FSHD primary muscle cells led to a significant downregulation of the pathway (Fig. S2A). Thus, DUX4 is required to activate a viral mimicry-like pathway in FSHD muscle cells.

**Fig. 2.**
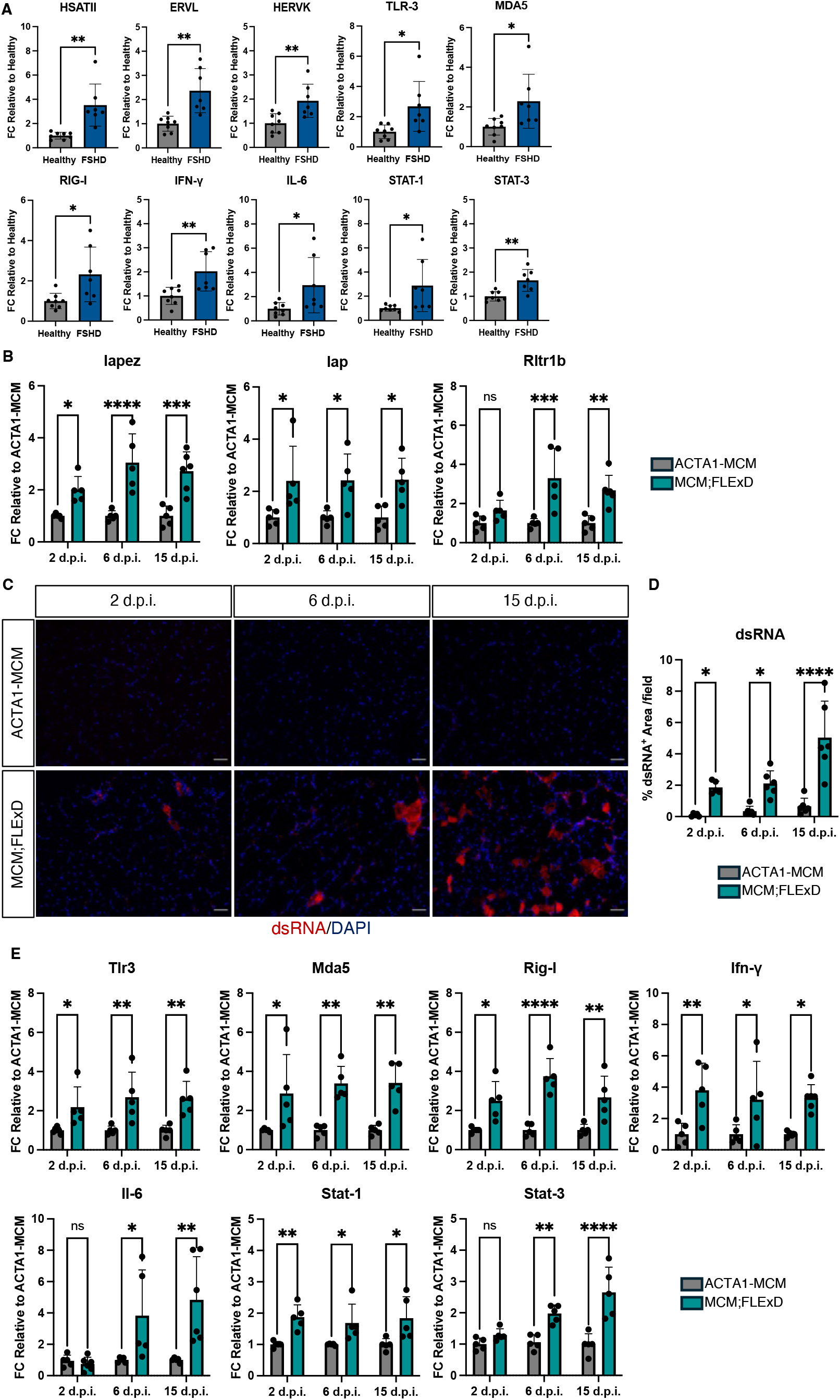
DUX4 activates a viral mimicry-like response. **(A)** Expression of genes encoding for repetitive elements targets of DUX4 (*HSATII*, *ERVL*, *HERVK*), dsRNA receptors (*TLR-3*, *MDA5*, *RIG-I*) and inflammatory factors (*IFN-γ*, *IL-6*, *STAT-1*, *STAT-3*) evaluated by RT-qPCR in differentiated human primary muscle cells from FSHD patients (n=7) compared to healthy donors (n=8). Values expressed as fold change relative to healthy donor cells. Statistical significance assessed by Unpaired t-test, *p<0.05; **p<0.01. **(B)** Expression of genes encoding for repetitive elements targets of DUX4 in mouse (*Iapez*, *Iap*, *Rltr1b*) evaluated by RT-qPCR on GA muscles from MCM;FLExD compared to ACTA1-MCM mice at 2-6-15 d.p.i.. n=5. Values expressed as fold change relative to ACTA1-MCM mice for each time point. Statistical significance assessed by two-way ANOVA, with Sidak’s multiple comparisons test, *p<0.05; **p<0.01; ***p<0.001; ****p<0.0001. **(C)** Representative images of dsRNA staining (red) on TA from ACTA1-MCM and MCM;FLExD mice at 2-6-15 d.p.i.. Nuclei stained with DAPI (blue). Scale bar: 50μm. **(D)** Quantification of the dsRNA signal in C. n=5-6. Statistical significance assessed by two-way ANOVA, with Sidak’s multiple comparisons test, *p<0.05; ****p<0.0001. **(E)** Expression of genes encoding for dsRNA receptors (*Tlr-3*, *Mda5*, *Rig-I*) and inflammatory factors (*Ifn-γ*, *Il-6*, *Stat-1*, *Stat-3*) evaluated by RT-qPCR in GA muscles from MCM;FLExD compared to ACTA1-MCM mice at 2-6-15 d.p.i.. n=5-6. Values expressed as fold change relative to ACTA1-MCM mice for each time point. Statistical significance assessed by two-way ANOVA, with Sidak’s multiple comparisons test, *p<0.05; **p<0.01; ****p<0.0001.

Next, we engineered human healthy donor muscle cells by introducing a doxycycline-inducible *DUX4* transgene (iDUX4) or its empty vector (iVector), as negative control. Interestingly, induction of DUX4 was sufficient to significantly activate its RE targets, leading to the accumulation of dsRNA and activation of the viral mimicry-like pathway in iDUX4 compared to iVector control cells (Fig. S2B-E). DUX4 induction was also sufficient to induce RE, cause dsRNA accumulation specifically in skeletal muscle fibers and significantly activate the viral mimicry-like pathway *in vivo* in MCM;FLExD compared to ACTA1-MCM mice (Fig. 2B-E and Fig. S3A-B). Hence, DUX4 is sufficient to activate a viral mimicry-like pathway in both human and mouse muscle cells.

In the cancer setting, activation of the viral mimicry pathway is usually associated to the induction of type I interferon (IFN-I), *IFN-α* and *IFN-β*, genes (46). Surprisingly, *IFN-I* genes were not significantly altered in FSHD or iDUX4 human muscle cells compared to their controls (Fig. S4A-B). In MCM;FLExD mice there was just a transient activation of *IFN*-*α* at six days post DUX4 induction, while *IFN*-*β* was never significantly altered (Fig. S4C). Thus, in human and mouse muscle, DUX4 activation induced specifically *IFN*-*γ*.

### Identification of the cell populations altered in MCM;FLExD mouse muscles

To determine the main cell types affected by the aberrant expression of DUX4, we compared tibialis anterior (TA) muscles from ACTA1-MCM and MCM;FLExD mice using single-nucleus RNA-seq (snRNA-seq). In ACTA1-MCM, snRNA-seq successfully identified all the principal cell types composing healthy mouse TA (Fig. 3A; Fig. S5A; Table S5). In MCM;FLExD, one of the most evident alterations was in the muscle fiber composition (Fig. 3A-B), in line with the fact that in MCM;FLExD DUX4 is expressed selectively by myofibers. While in ACTA1-MCM type 2A, type 2B and type 2X muscle fibers constituted large clusters, they were strongly depleted in MCM;FLExD mice and were substituted by a cluster that we named “DUX4-induced aberrant myofiber” (Fig. 3C; Fig. S5B). Indeed, this was the only cluster expressing the DUX4 signature target genes (Table S6 and Methods) (Fig. 3D).

**Fig. 3.**
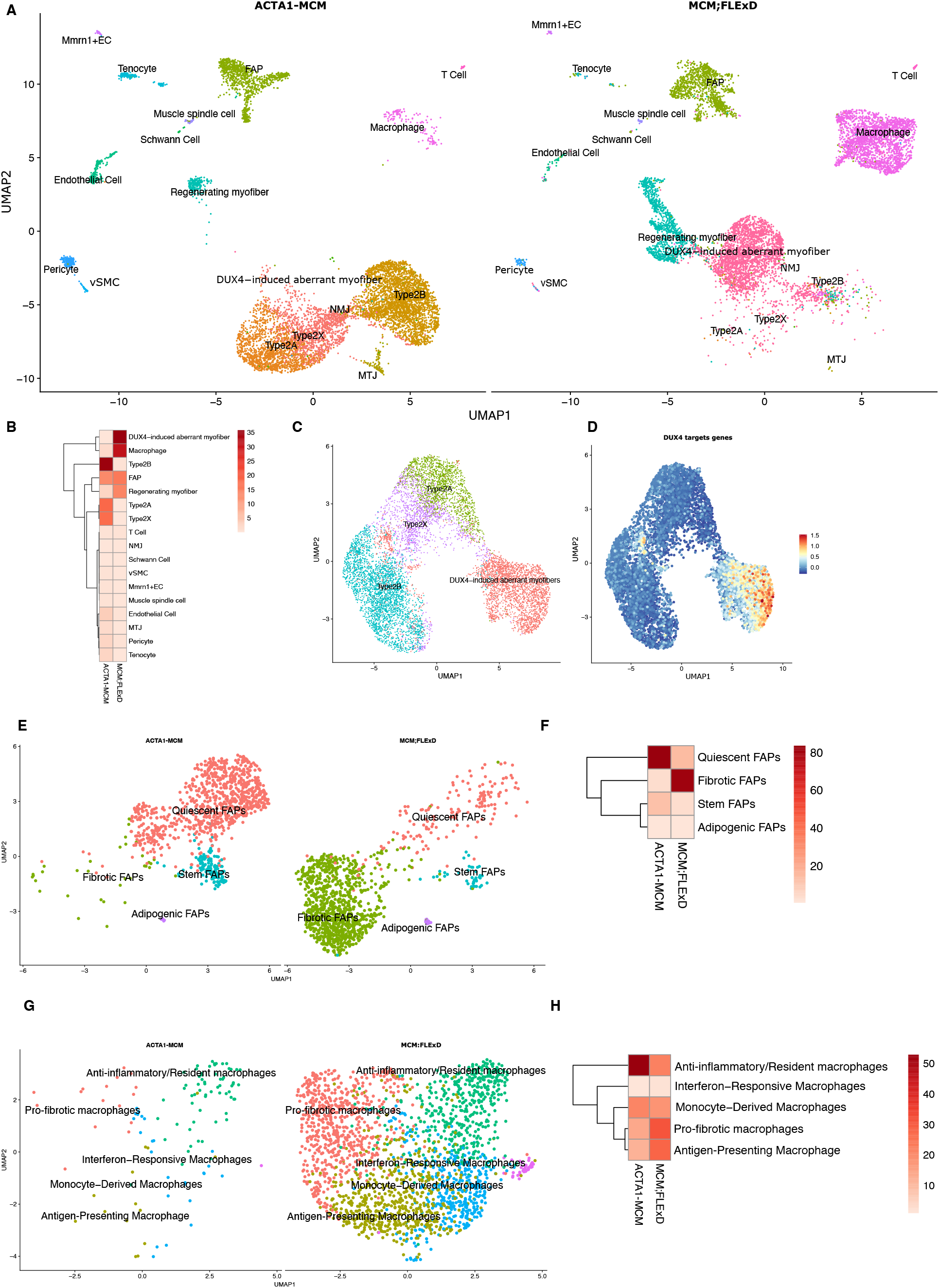
Single-nucleus RNA-seq identifies altered cell populations in MCM;FLExD muscle. **(A)** UMAP embedding of snRNA-seq data colored by cluster in ACTA1-MCM and MCM;FLExD. **(B)** Heatmap showing the frequency (%) of cell populations in A. **(C)** UMAP embedding colored by myofiber clusters in ACTA1-MCM and MCM;FLExD. **(D)** UMAP embedding colored by DUX4 signature expression in ACTA1-MCM and MCM;FLExD, confirming restricted expression in DUX4-induced aberrant myofibers. **(E)** UMAP embedding of FAPs colored by sub-cluster in ACTA1-MCM and MCM;FLExD. **(F)** Heatmap showing the frequency (%) of FAPs sub-clusters in E. **(G)** UMAP embedding of macrophages colored by sub-cluster in ACTA1-MCM and MCM;FLExD. **(H)** Heatmap showing the frequency (%) of macrophage sub-clusters in G.

Beside muscle fibers, another cluster affected by DUX4 expression was the one of fibro-adipogenic progenitors (FAPs). FAPs are a heterogenous mesenchymal cell population, interacting with muscle stem cells and immune cells, which is important to support skeletal muscle repair and homeostasis, but driving fibrosis and fat accumulation in disease (48,49). Compared to ACTA1-MCM, in MCM;FLExD FAPs were increased (Fig. 3B) and displayed an evident alteration of their subsets characterized by a decrease in quiescent- and stem-like FAPs and an expansion of pro-fibrotic FAPs (Fig. 3E-F; Fig. S5C).

An additional cell population showing a marked alteration in MCM;FLExD compared to ACTA1-MCM was macrophages, multifunctional cells that are critical for muscle homeostasis and disease pathogenesis (50). Here, we discovered both quantitative and qualitative alterations. Indeed, despite all macrophages populations were generally increased in MCM;FLExD compared to ACTA1-MCM mice (Fig. 3B), we also observed a relative reduction in anti-inflammatory/resident macrophages (51) and a relative expansion in pro-fibrotic and antigen presenting macrophages (51,52) (Fig. 3G-H; Fig. S5D).

We validated the alteration in the abundance of the main cell types by monitoring the expression level of cell type specific markers (Fig. S6A).

Collectively, our results indicate that muscle fibers, FAPs and macrophages are the main cell populations affected by aberrant DUX4 expression.

### Spatial mapping of muscle tissue degeneration in MCM;FLExD mice

As a complementary approach to snRNA-seq and to characterize the spatial organization of the muscle tissue in MCM;FLExD compared to ACTA1-MCM mice, we performed spatial transcriptomics (ST) using the 10x Genomics Visium HD platform. Specifically, we conducted ST on TA muscles from ACTA1-MCM and MCM;FLExD mice at multiple time points upon DUX4 induction (2, 6, and 15 d.p.i.). By integrating transcriptomic data with histological context, we delineated the spatial distribution of gene expression and cell populations across tissue sections while preserving the spatial context. Notably, all major cell populations identified by snRNA-seq were also recapitulated in the spatial dataset (Fig. 4A-B; Table S7). Consistent with our single-nucleus analysis, one of the most prominent alteration in MCM;FLExD mice was in muscle fiber composition, with a progressive expansion of the DUX4-induced aberrant myofibers (Fig. 4C-D; Fig. S7A) and a progressive reduction of Type 2A/2B/2X myofibers at the 2, 6, and 15 d.p.i. timepoints in MCM;FLExD compared to ACTA1-MCM mice (Fig. 4C; Fig. S7A). Spatially speaking, the first area showing the appearance of DUX4-induced aberrant myofibers, already two days after DUX4 induction, was the one occupied by Type 2B myofibers. Instead, DUX4-induced aberrant myofibers appeared in the Type 2A/2X myofiber occupied area only starting from six days after DUX4 induction (Fig. S7A). Unlike snRNA-seq, ST allowed to identify also Type 1 myofibers, which were a minor population in TA (Fig. 4A-B), as expected, and were not dramatically altered by DUX4 expression (Fig. 4C, Fig. S7A). Finally, MCM;FLExD mice showed an increase in the FAPs, macrophages and regenerating myofiber populations across the entire muscle section area (Fig. 4C; Fig. S7A).

**Fig. 4.**
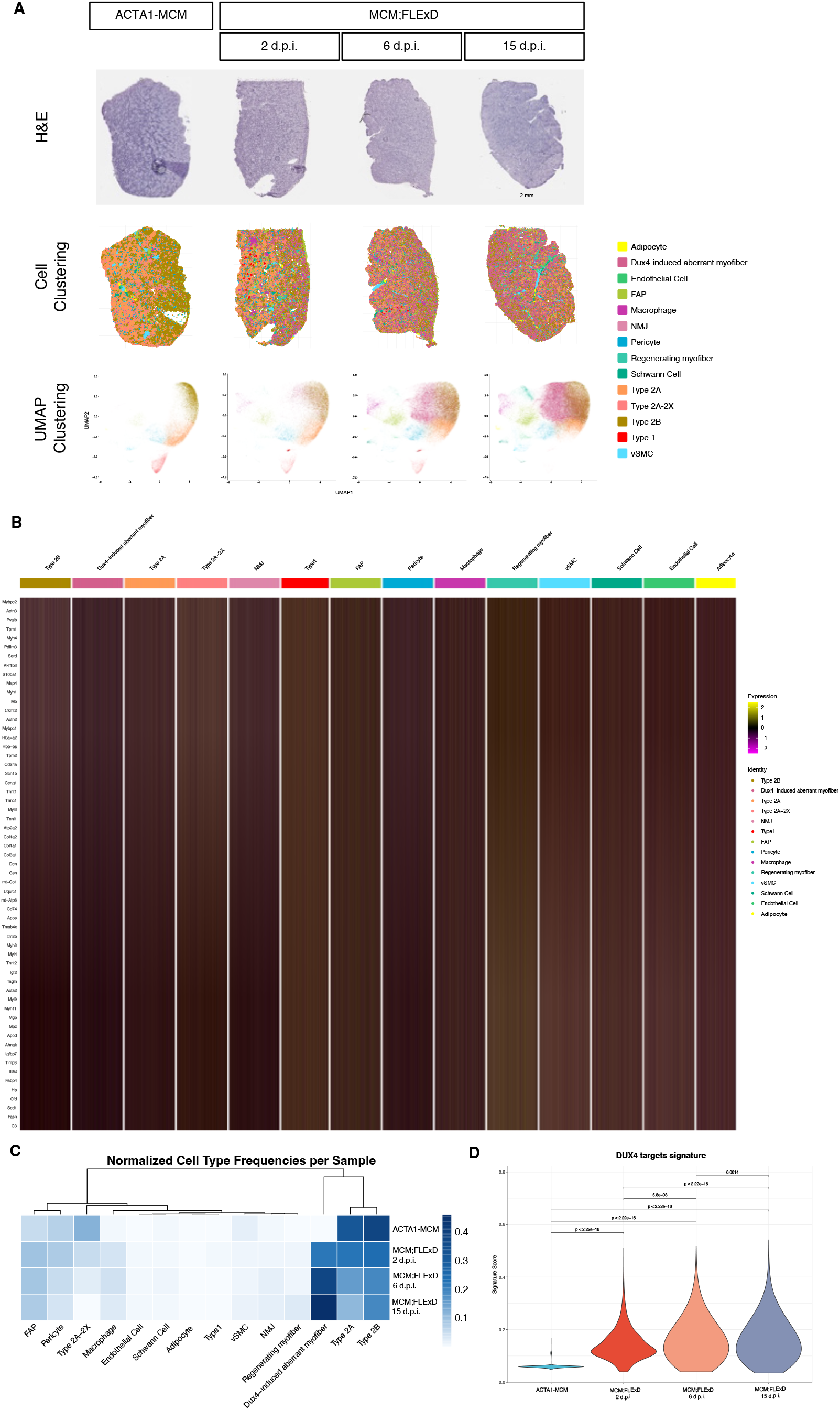
Spatial Transcriptomics profiling of ACTA1-MCM and MCM;FLExD muscles. **(A)** H&E-stained (top), cell clusters overlaid on the tissue-covered area (center) and UMAP plots (bottom) of TA muscle sections from ACTA1-MCM and MCM;FLExD mice at 2-6-15 d.p.i.. **(B)** Heatmap for the top enriched genes per cluster in ACTA1-MCM and MCM;FLExD. **(C)** Heatmap showing the frequency of the cell populations in ACTA1-MCM and MCM;FLExD at 2-6-15 d.p.i.. **(D)** Violin plots showing DUX4 targets UCell signature scores across ACTA1-MCM and MCM;FLExD at 2-6-15 d.p.i. myofibers. Statistical significance brackets show pairwise Wilcoxon rank-sum test results.

### IFN-γ expression is restricted to MCM;FLExD muscle fibers

According to our snRNA-seq and ST results, the DUX4-induced aberrant myofibers were the main ones expressing the IFN-γ response gene set (Fig. 5A-B). Importantly, expression of *IFN-γ* (Fig. 2E) and activation of the IFN-γ response pathway (Fig. 5B) were already evident at 2 d.p.i., when muscle degeneration is minimal (Fig. S1B) and (30).

**Fig. 5.**
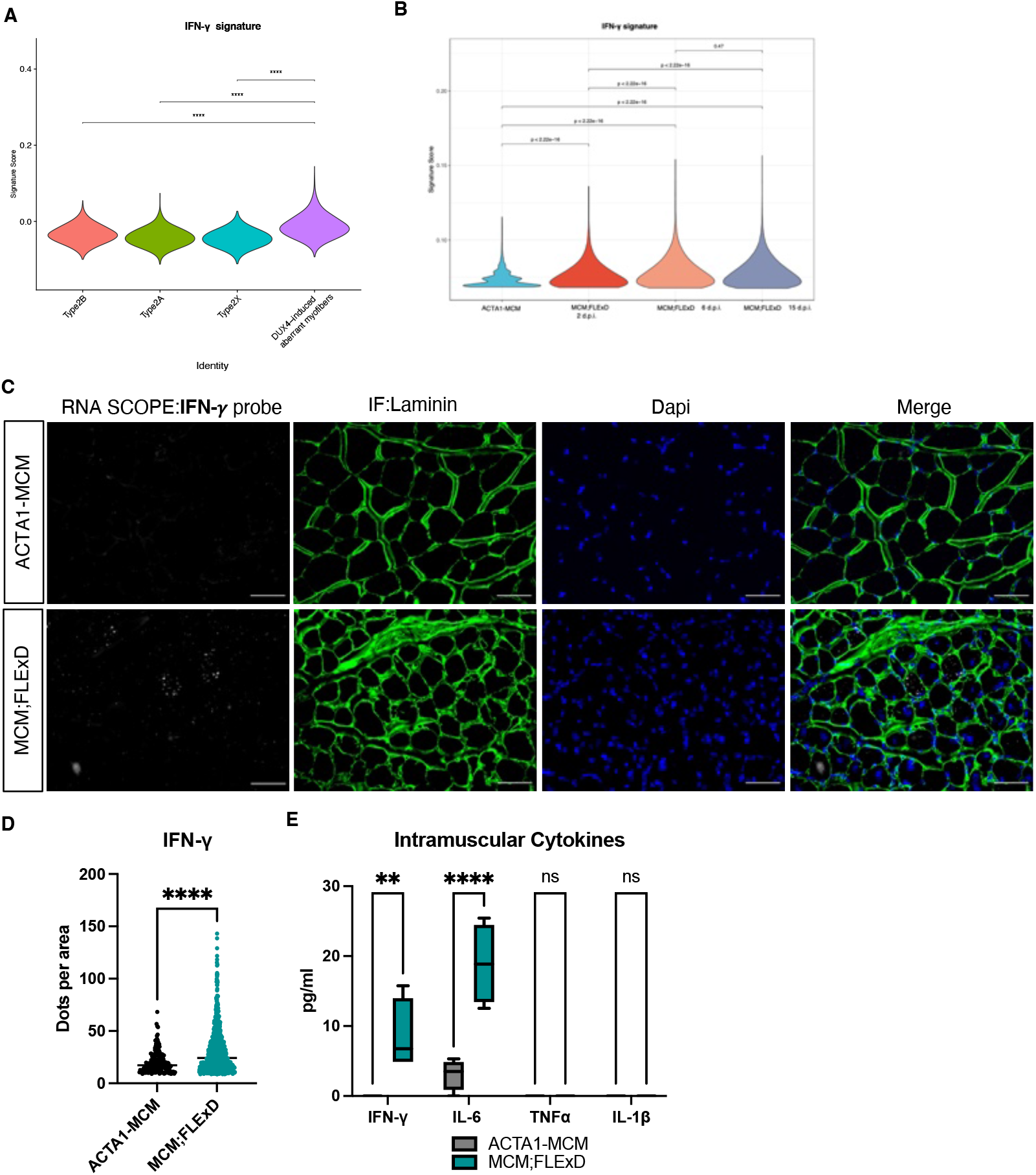
IFN-γ is produced specifically by MCM;FLExD myofibers. **(A)** Violin plots showing IFN-γ signature scores in ACTA1-MCM and MCM;FLExD myofiber types based on snRNAseq data. Statistical significance brackets show pairwise Wilcoxon rank-sum test results. **(B)** Violin plots showing IFN-γ UCell signature scores in myofibers of ACTA1-MCM and MCM;FLExD at 2-6-15 d.p.i. based on ST data. Statistical significance brackets show pairwise Wilcoxon rank-sum test results. **(C)** Representative images of RNAscope multiplex fluorescent assays detecting *Ifn-γ* RNA (grey) combined with immunofluorescence for laminin (green), nuclear staining with DAPI (blue), in GA from ACTA1-MCM and MCM;FLExD mice at 15 d.p.i.. **(D)** Quantification of the *Ifn-γ* dots per area in C. Statistical significance assessed by Unpaired t-test, ****p<0.0001. **(E)** Luminex Assays on GA protein extracts from ACTA1-MCM and MCM;FLExD mice at 6 d.p.i.. n=4. Statistical significance assessed by Two-way analysis of variance (ANOVA) with Sidak’s multiple comparisons test, **p<0.01; ****p<0.0001.

To identify the source of *IFN-γ* expression, we performed RNAscope with a probe specific for *IFN-γ* in combination with immunofluorescence for laminin. Remarkably, we found that *IFN-γ* expression is restricted to MCM;FLExD myofibers (Fig. 5C-D).

To validate the expression of IFN-γ at protein level, we performed Luminex assays on skeletal muscle extracts of ACTA1-MCM and MCM;FLExD mice. Together with IL-6, which we used as positive control since it was previously found upregulated in FSHD patients and MCM;FLExD mice (43,53,54), we found significantly increased IFN-γ levels in MCM;FLExD compared to ACTA1-MCM mice (Fig. 5E). Notably, intra-muscular levels of TNF-α and IL-1β were not significantly altered in MCM;FLExD compared to ACTA1-MCM mice (Fig. 5E) supporting a selective alteration of pro-inflammatory cytokines in MCM;FLExD myofibers.

Since there are multiple reports of FAP alteration in FSHD patients and animal models (31,55–60), we analyzed them in more detail. We found a significant increase in the FAPs marker *PDGFRα* in MCM;FLExD compared to ACTA1-MCM mice at all timepoints analyzed (Fig. S8A). FACS analysis confirmed the significant expansion of the FAP population in MCM;FLExD compared to ACTA1-MCM mice (Fig. S8B-C). Moreover, starting from day 6 d.p.i., we found a significant accumulation of fibrotic and adipogenic infiltrate in MCM;FLExD compared to ACTA1-MCM muscle (Fig. S8D-G). Next, we purified FAPs from ACTA1-MCM and MCM;FLExD muscles by FACS sorting and monitored *IFN-γ* and *IL-6* expression. While no *IFN-γ* expression was detectable, FAPs purified from MCM;FLExD muscle expressed significantly higher *IL-6* compared to control mice (Fig. S8H). Collectively, these results support the conclusion that IFN-γ is selectively produced by MCM;FLExD muscle fibers.

### IFN-γ is sufficient to recapitulate several DUX4 effects

Beside DUX4-induced aberrant myofibers, the IFN-γ response gene set was activated also in regenerating myofibers, FAPs and macrophages of MCM;FLExD mice (Fig. 6A-B), suggesting that IFN-γ produced by MCM;FLExD myofibers has a broad effect. To test directly its relevance, we treated human healthy donor primary muscle cells *in vitro* with recombinant IFN-γ. We also injected recombinant IFN-γ *in vivo* intramuscularly in wild-type (WT) mice. Notably, in both settings we found that IFN-γ was sufficient to activate the expression of dsRNA receptors and pro-inflammatory transcription factors (Fig. 6C-D). IFN-γ also had a substantial functional impact. When administered to actively growing healthy donor primary muscle cells, IFN-γ significantly inhibited their proliferation compared to vehicle-treated cells (Fig. 6E; G). On the other end, IFN-γ significantly inhibited myotube formation if added to the cells after they have reached confluency and have been shifted to muscle differentiation (Fig. 6F; H). *In vivo*, IFN-γ was sufficient to significantly increase the number of intramuscular FAPs and macrophages (Fig. 6I-L). Thus, IFN-γ is sufficient to recapitulate several pathogenic effects associated to aberrant DUX4 expression.

**Fig. 6.**
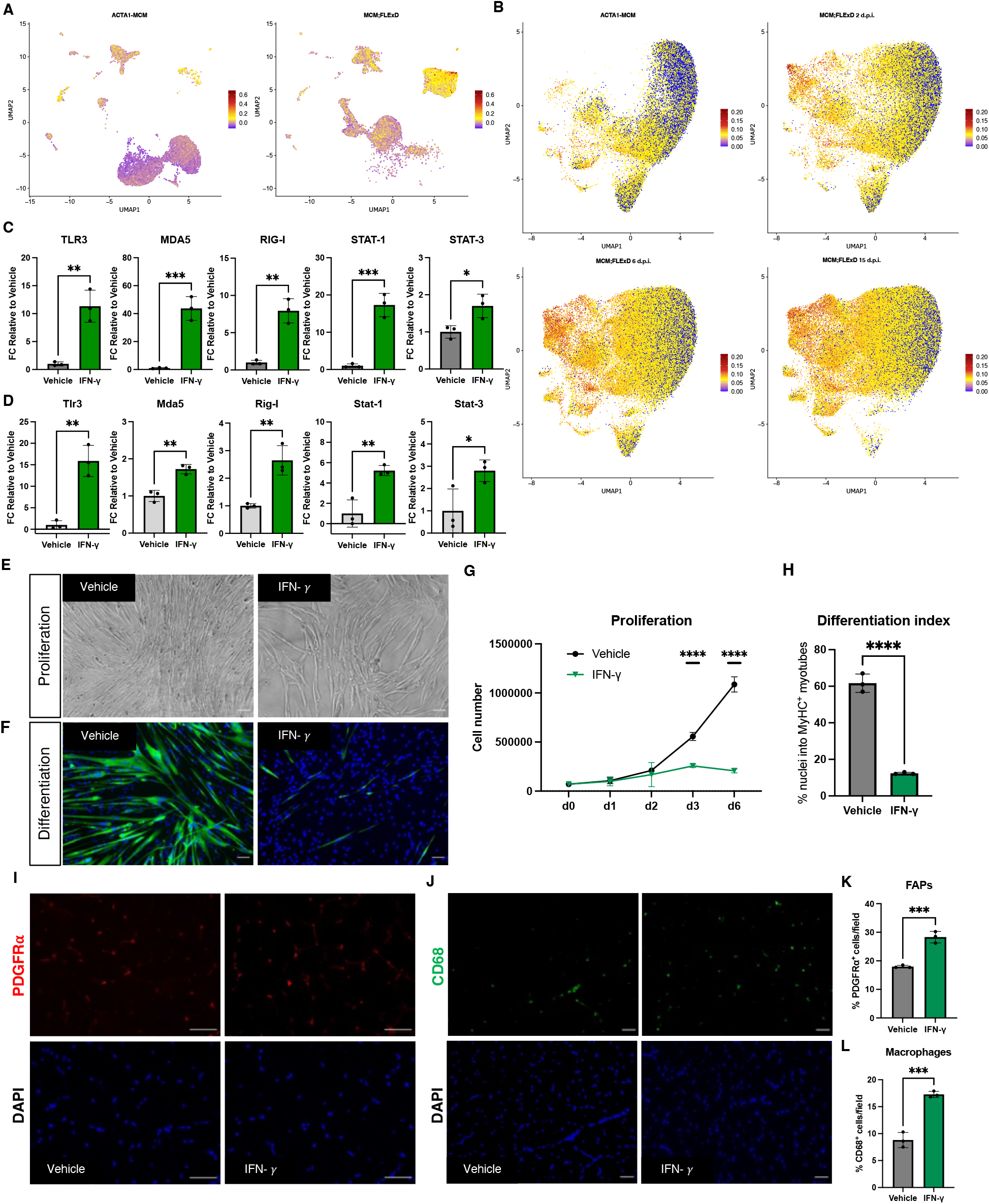
IFN-γ is sufficient to recapitulate several DUX4 effects. **(A)** UMAP embedding of snRNA-seq data colored by IFN-γ signature expression in ACTA1-MCM and MCM;FLExD muscles. **(B)** UMAP embedding of ST data colored by IFN-γ signature expression in ACTA1-MCM and MCM;FLExD at 2-6-15 d.p.i.. **(C)** Expression of the indicated viral mimicry genes evaluated by RT-qPCR in human healthy donor muscle cells treated with vehicle or IFN-γ. Values expressed as fold change relative to vehicle treated cells. n=3. Statistical significance assessed by Unpaired t-test, *p<0.05; **p<0.01; ***p<0.001. **(D)** Expression of the indicated viral mimicry genes evaluated by RT-qPCR in GA of wt mice treated with vehicle or IFN-γ. n=3. Statistical significance assessed by Unpaired t-test, *p<0.05; **p<0.01. **(E)** Representative phase contrast images of human healthy donor muscle cells treated with vehicle or IFN-γ during proliferation. Scale bar: 50μm. **(F)** Representative images of immunofluorescence for MHC (MF20, green), nuclear staining with DAPI (blue) of human healthy donor muscle cells treated with vehicle or IFN-γ during myogenic differentiation. Scale bar: 50μm. **(G)** Quantification of results in E. Curves represent the means ± SD. n=3. Statistical significance assessed by Two-way analysis of variance (ANOVA), with Sidak’s multiple comparisons test, ****p<0.0001. **(H)** Quantification of results in F. Statistical significance assessed by Unpaired t-test, ****p<0.0001. **(I)** Representative images of GA muscle from wt mice treated with vehicle or IFN-γ stained for PDGFR*α* (red), nuclear staining with DAPI (blue). Scale bar: 50μm. **(J)** Representative images of GA muscle from wt mice treated with vehicle or IFN-γ stained for CD68 (green), nuclear staining with DAPI (blue). Scale bar: 50μm. **(K)** Quantification of results in I. n=3. **(L)** Quantification of results in J. n=3. For K and L statistical significance assessed by Unpaired t-test, ***p<0.001.

### IFN-γ is required for disease progression downstream of DUX4

Our data position IFN-γ immediately downstream of DUX4 making it a strong candidate to mediate its pathogenic effects. To test this hypothesis, we treated MCM;FLExD mice with an anti-IFN-γ neutralizing monoclonal antibody (IFN-γ NAb) or its isotype control (IgG-Ctrl) (Fig. 7A). Notably, we found that IFN-γ NAb treatment led to an evident amelioration of the muscle histological features compared to IgG-Ctrl (Fig. 7B). To quantify the effect, we performed specific staining for FAPs, macrophages, muscle fibro-fatty infiltration, muscle atrophy and necrosis (Fig. 7C-E and Fig. S9A). Compared to MCM;FLExD mice treated with IgG-Ctrl, IFN-γ neutralization led to a significant reduction in FAP and macrophage expansion associated to a significant decrease in muscle fibro-fatty infiltration (Fig. 7F-J). This was associated to a significant preservation of muscle fiber Cross-Sectional Area (CSA), a key indicator of muscle atrophy, in MCM;FLExD mice treated with IFN-γ NAb compared to the IgG-Ctrl (Fig. 7K and S9B-C). In line with the fact that the IFN-γ neutralization does not block the expression of DUX4 (Fig. S9D), we found no significant change in the number of necrotic fibers (Fig. S9E).

**Fig. 7.**
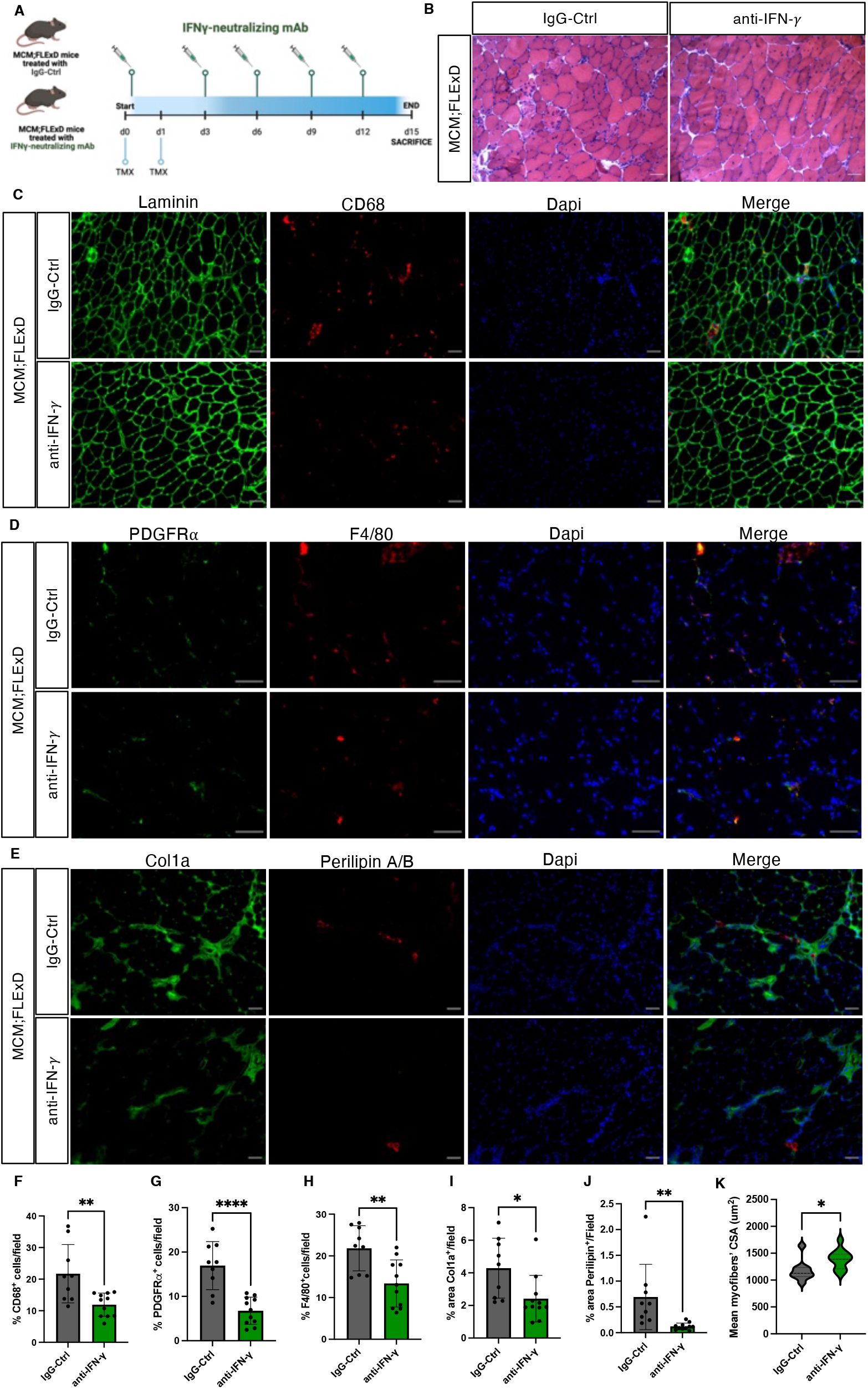
IFN-γ neutralization reduces dystrophic features of MCM;FLExD mice. **(A)** Schematic representation of the experimental approach. Created with BioRender.com. **(B)** Representative images of H&E staining of TA from MCM;FLExD mice treated with control IgG (IgG-Ctrl) or anti-IFN-γ NAb. Scale bar: 50μm. **(C)** Representative images of immunofluorescence for laminin (green), CD68 (red) and nuclei in DAPI (blue) on TA from MCM;FLExD mice treated with IgG-Ctrl or anti-IFN-γ NAb. Scale bar: 50μm. **(D)** Representative images of immunofluorescence for PDGFR*α* (green), F4/80 (red) and nuclei in DAPI (blue) on TA from MCM;FLExD mice treated with IgG-Ctrl or anti-IFN-γ NAb. Scale bar: 50μm. **(E)** Representative images of immunofluorescence for Col1a (green), Perilipin (red) and nuclei in DAPI (blue) on TA from MCM;FLExD mice treated with IgG-Ctrl or anti-IFN-γ NAb. Scale bar: 50μm. **(F-J)** Quantification of the results show in C, D, E. The graph represents the percentage of positive cells per field. (n=9 for IgG-Ctrl, n=11 for anti-IFN-γ). **(K)** Violin plot quantification of the CSA for the results in C (n=9 for IgG-Ctrl, n=11 for anti-IFN-γ). Statistical significance assessed by Unpaired t-test, *p<0.05; **p<0.01; ****p<0.0001.

Collectively, our results indicate that IFN-γ is a critical driver of disease progression downstream of DUX4.

## DISCUSSION

Despite the inappropriate gain of DUX4 expression was identified as the cause of FSHD long ago (61), the molecular pathogenesis of the disease is still unclear, and no cure or therapeutic option is available to FSHD patients. Currently, most therapeutic efforts are devoted on developing ways to block DUX4 expression [reviewed in (62)]. Nevertheless, there are multiple reports that DUX4 downstream pathways might persist upon its silencing (3–11). Therefore, the identification of DUX4-activated pathways required for the pathogenesis of FSHD bears strong therapeutic relevance. Among them is muscle inflammation which, in FSHD, displays uncommon features. In other muscular dystrophies, inflammation is usually secondary to muscle degeneration [reviewed in (13)]. Instead, inflammation is an early sign in FSHD, occurring before muscle atrophy and its fibro-fatty substitution (12). FSHD muscle cells have been shown to activate an innate immune inflammatory response, which leads to an immune-mediated muscle loss (25). Accordingly, muscle inflammation in FSHD is associated to active disease (12,28,29,31,63–67), and is increasingly recognized as driver of disease progression (12,14–16). Nevertheless, the molecular mechanism of FSHD muscle inflammation remains poorly understood and no effective anti-inflammatory treatment has been identified yet (17–21).

To decipher muscle inflammation in FSHD, we used an unbiased approach. By analyzing publicly available gene expression profiling from six independent studies, totaling to 59 healthy donor and 218 FSHD patient muscle biopsies, we consistently identified “IFN-γ response” as the most significantly upregulated inflammatory pathway in FSHD. We obtained the same result in the MCM;FLExD mouse model of the disease. Intriguingly, we found that, within FSHD patients, the IFN-γ response pathway is particularly upregulated in inflamed compared to not inflamed muscles, even in the very same patient.

We found that treatment with recombinant IFN-γ is sufficient to inhibit myogenic differentiation, and activate both FAPs and macrophages, as already described for DUX4 (58,60,68–70). Previous work also showed that IFN-γ activates the complement system, mitochondrial dysfunction, oxidative stress, and ferroptosis, which have all been described in FSHD (27,28,71–101). Hence, IFN-γ activation is associated to many aspects of the FSHD pathogenesis.

Recently, a role for IFN-γ in idiopathic immune myopathies was reported (102). While in that setting the source of IFN-γ are CD8 T cells, in line with our results recombinant IFN-γ treatment reduced both proliferation and fusion of human muscle cells. Chronic exposure to recombinant IFN-γ of WT mice subjected to acute sterile muscle injury significantly impaired muscle repair leading to muscle atrophy, while promoting fibrosis and fat deposition (102).

Collectively, these results strongly suggest that IFN-γ produced by muscle fibers actively contributes to FSHD disease progression.

Through single-nucleus and spatial transcriptomics profiling, we generated a comprehensive atlas offering unprecedented resolution of the dystrophic tissue microenvironment for studying FSHD disease biology that can be queried to explore how local cell–cell communication contributes to dystrophic disease progression, and for biomarker and therapeutic target discovery.

A previous proteomics study found a significant increase of intramuscular IFN-γ in inflamed compared to not inflamed muscles from the very same FSHD patient (103). Intriguingly, we found that IFN-γ is directly produced by FSHD muscle cells and MCM;FLExD muscle fibers. Despite *IFN-γ* is not a direct DUX4 target (104–108), its expression is intimately linked to that of DUX4. We found that, in human muscle cells, DUX4 is required and sufficient to activate *IFN-γ*. Contrary to *IL-6*, in MCM;FLExD mice *IFN-γ* is significantly increased already two days after DUX4 activation, when we and others (30) can detect a negligent level of muscle degeneration. Therefore, the production of IFN-γ does not appear just one of many downstream byproducts of muscle fiber death, as it is the case in other muscular dystrophies [reviewed in (13)].

What could be the mechanism driving *IFN-γ* activation in FSHD? We found activation of a viral mimicry-like pathway in muscle cells of FSHD patients and in MCM;FLExD muscle. Nevertheless, we surprising found no *IFN-I* gene activation. At first glance, this appears paradoxical given the robust accumulation of endogenous dsRNA and induction of pattern recognition receptors. However, the available data suggest that this reflects an active uncoupling of viral sensing from *IFN-I* transcription rather than a defect in innate immune recognition.

The canonical viral mimicry pathway starts with the accumulation of dsRNA and its recognition by dsRNA receptors, such MDA5, RIG-I or TLR3, which signal through TANK-binding kinase 1 (TBK1) and TGF-beta-activated kinase 1 (TAK1). TBK1 activates the interferon regulatory transcription factor 3 (IRF3), which induces the expression of *IFN-I* genes (109). Instead, TAK1 activates the transcription factors NF-κB, CREB, and AP1 that cooperates to promote the induction of several pro-inflammatory cytokines, including IFN-γ (110). Thus, the dsRNA-induced pathway splits downstream of dsRNA sensing: the IRF3-dependent arm leads to IFN-I induction, while the stress-responsive arm activates inflammatory cytokines.

Among the genes upregulated by DUX4 in FSHD (111), there are multiple known *IFN-I* gene repressors. For example, PPM1B (Protein Phosphatase, Mg2+/Mn2+ Dependent 1B) dephosphorylates and inactivates TBK1 and IRF3, thereby suppressing *IFN-I* gene expression (112). In addition, the phosphate transporter SLC34A2 (solute carrier family 34 member 2) inhibits TBK1 kinase activity, preventing *IFN-I* expression (113). Instead, among the genes downregulated by DUX4 in FSHD (111), there are several essential activators of IFN-I. For example, the DEXD/H box helicase DDX60 is required for *IFN-I* gene activation downstream of RIG-I and MDA5 (114). Also, TRIM56 (Tripartite Motif-containing 56) is required for activation of IRF3 and *IFN-I* gene induction (115). Intriguingly, DUX4 has been shown to decrease the phosphorylation of TBK1 and IRF3 and suppress *IFN-I* gene activation following herpesvirus infection (113). Based on these results, it is tempting to speculate that, despite innate immune activation, DUX4 might antagonize a pathway essential for IFN-I but not strictly required for IFN-γ production (116,117). This distinction is critical for understanding how DUX4 might drive chronic inflammation without resolution in FSHD muscle. Unlike classical viral mimicry, in which endogenous dsRNA elicits a productive IFN-I response leading to immune clearance, in FSHD DUX4 appears to induce a state of “skewed viral mimicry” in which innate immune sensors are activated but selectively uncoupled from interferon-mediated antiviral resolution. These uncoupling favors persistent IFN-γ–dominated inflammation, tissue damage, and disease progression.

IFN-γ binding to its receptor initiates a cascade of events wherein the associated Janus kinases, JAK1 and JAK2, activate several transcription factors including STAT1, STAT3, CIITA, NF-κB, AP-1, EGR1, EGR2, C/EBP-β, ATF2 and ATF6, which subsequently induce interferon-stimulated genes (ISGs) [reviewed in (118)]. Since DUX4 has been shown to repress IFN-γ signaling by inhibiting STAT1 (119), it is conceivable that the activation of the IFN-γ response pathway that we identified takes place via the IFN-dependent, STAT1-independent pathway (120–131).

Future studies aiming to dissect the exact mechanism through which DUX4 regulates the IFN pathway in skeletal muscle may offer additional strategies by which muscle inflammation may be targeted in FSHD.

While we found that IFN-γ NAb blocks the progression of the disease in the MCM;FLExD model of FSHD, the main limitations of this study are that we treated pre-symptomatic mice and we did not validate the long-term safety of IFN-γ inhibition. In the future, it would be important to neutralize IFN-γ in symptomatic mice, as well as to assess the long-term tolerability of the treatment.

When assessed by the treadmill running test, MCM;FLExD have been reported to show a significant functional deficit compared to ACTA1-MCM mice (132). Despite the significant histological amelioration caused by IFN-γ NAb is expected to rescue muscle function, we decided not to assess this aspect upon IFN-γ neutralization. Indeed, IFN-γ and IL-6 are known to be modulated by physical activity (133–136), which would have been a strong confounding factor for our study. Future functional studies, possibly not involving physical activity, are thus required to determine if IFN-γ neutralization rescues also MCM;FLExD muscle strength.

Inhibitors of the IFN-γ pathway are approved for several autoimmune and inflammatory diseases and are currently in phase 2 or 3 clinical trials for several other inflammatory conditions (137,138). Future assessment of safety, tolerability, and toxicity of IFN-γ pathway inhibitors in pre-clinical models will help determine the therapeutic potential of IFN-γ inhibition for FSHD.

## MATERIALS AND METHODS

### Animals

Mouse studies were approved by the Institutional Animal Care and Use Committee of San Raffaele and by the Italian Ministry of Health (Rome, Italy) (IACUC approval 607/2023-PR), and were performed in full compliance with relevant ethical regulations. FLExDUX4 [B6(Cg)-Gt(ROSA)26Sortm1.1(DUX4*)Plj/J (JAX 028710)] and ACTA1-MCM [Tg(ACTA1-cre/Esr1*)2Kesr/J (JAX 025750)] were purchased from The Jackson Laboratory (Maine, USA). WT C57BL/6N mice (027C57BL/6) were purchased from Charles River Laboratories (Calco, Italy). Animals were bred in-house and housed in a temperature and humidity-controlled room in a specified pathogen-free environment under 12:12 h light/dark cycles. 10-12 weeks-old mice were used for all experiments. Animals were weighed at the beginning of the study and randomly allocated to treatment groups to ensure similar mean body weight across groups.

### Mouse genotyping

DNA was isolated from 2 mm tail snips from <10-day-old mice. DNA extraction and PCR were performed with Phire Tissue Direct PCR Master Mix (Thermo Scientific #F-170S) following the manufacturer’s instructions, and using the primers reported in The Jackson Laboratory instructions for the mouse strains (Table S8).

### Tamoxifen administration

Tamoxifen (TMX) administration was performed as in (42). Specifically, TMX (Sigma-Aldrich #T5648) was dissolved in 100% ethanol at 55 °C and added to warm sterile corn oil to make a 20 mg/ml stock. Stock TMX aliquots were briefly warmed to 37 °C and diluted further with warm corn oil just prior to use. Animals were weighed and injected IP on two consecutive days (d0 and d1) with the appropriate volume of TMX for a final concentration of 5 mg/kg.

### Isolation of primary muscle cells from healthy donors and FSHD patients

Human sample collection was approved by the Institutional Ethics Committees of Koç University (Protocol code 2020.298.IRB1.106) and Azienda Ospedaliera Universitaria Pisana (Protocol code EJPRD). Informed consent was obtained from all participants. Participants gave blood samples for molecular testing and confirmation of the disease status at the time of clinical evaluation. Bilateral thigh MRIs were obtained to determine the status of the donor muscles. Before surgery, a re-evaluation for clinical scoring was performed to confirm that the patient had not progressed to a more severe phase of the disease. Next, an MRI (single thigh) with markings was obtained on the same day. Coronal images were used to determine the biopsy level in relation to the closest marking capsule. Needle muscle biopsies and primary myoblast cell cultures were performed as indicated by the FSHD Research Center at University of Rochester Medicine (https://www.urmc.rochester.edu/neurology/fshd-center/research-info/protocols). Next, muscle cells were purified using CD56 MicroBeads (Miltenyi Biotech #130-050-401). Evaluation of muscle cell purity was performed by staining with anti-desmin antibody (Leica #DES-DERII-L-CE) followed by FACS quantification of the percentage of desmin positive cells.

### Primary cell culture and differentiation

Use of human muscle cells for research was approved the Institutional Ethics Committee of IRCCS Ospedale San Raffaele (Protocol code FSHD). Information on human primary muscle cells derived from FSHD patients and healthy donors is available in Table S4. Cells were cultured and maintained at 37°C in a humidified incubator with 5% O_2_ and 5% CO_2_. Myoblasts were grown in Ham’s F-10 medium (Euroclone, #ECB7503L), supplemented with L-Glutamine (Thermo Fisher Scientific, #25030081), 20% FBS (Thermo Fisher Scientific, #A5256701), 1% Pen/Strep (Thermo Fisher Scientific, #15140130), 10 ng/ml bFGF (Tebu-bio, #100-18B), and 1 μM dexamethasone (Sigma-Aldrich, #D4902). To induce myogenic differentiation, growth medium was replaced by DMEM:F12 (1:1, Sigma-Aldrich, #D9785) supplemented with 20% knockout serum replacement (KOSR, Thermo Fisher Scientific, #10828028), 3.151 g/L glucose, 10 mM MEM non-essential amino acids (Thermo Fisher Scientific, #11140050), 1 mM sodium pyruvate (Thermo Fisher Scientific, #11360070). Differentiation was carried out for 96h.

### Establishment of muscle cells with doxycycline-inducible DUX4 (iDUX4)

Immortalized healthy donor human muscle cells (MB161; 01Ubic Ct5) were obtained from the UMass Chan Wellstone Center for FSHD (https://www.umassmed.edu/wellstone/about/overview/). Cells were plated on gelatin (0.1% in water, Sigma #G9391)-coated dishes in LHCN Medium (DMEM (Euroclone #ECM0728L), Medium 199 1X (Gibco #31150-022), 15% Tetracycline Negative FBS (Euroclone #ECS0182L), 1% Pen/Strep, 1.4 mg/L vitamin B12 (Sigma #V2876), 0.03 mg/l ZnSO4 (Fisher Scientific Z68-500), 0.02M Hepes buffer (Sigma-Aldrich #H3375), 10 μg/L bFGF (Tebu-bio, #100-18B), 2.5 μg/L HGF (Tebu-bio, #100-39), 0.055 mg/L dexamethasone (Sigma-Aldrich, #D4902). Cells were cultured at 37°C in a humidified incubator with 5% O_2_ and 5% CO_2_.

Lentiviruses encoding doxycycline-inducible wild-type DUX4 (107) and the corresponding control vector pCW57.1 were produced in HEK293T cells using the calcium phosphate transfection method. To establish stable cell lines, cells were plated at low density and transduced with lentiviruses in the presence of 8 µg/mL polybrene (Sigma-Aldrich #TR-1003-G). Transduced cells were selected with 0.5 µg/mL puromycin (Sigma-Aldrich, #P9620). Following selection, doxycycline-inducible DUX4 (iDUX4) or empty vector (iVector) cells were maintained under continuous puromycin selection.

### Recombinant IFN-γ treatments

To evaluate its effect on cell growth, proliferating healthy donor primary human muscle cells were plated in 12-well plates at a density of 70.000 cells/well in growth media. Recombinant Human IFN-γ (R&D Systems #285-IF) was added at a concentration of 25 ng/ml. PBS was used as vehicle. The medium with PBS or IFN-γ was refreshed daily.

To assess its effect on myogenic differentiation, healthy donor primary human muscle cells were plated in 6-well plates at a density of 460.000 cells/well in growth media. The following day, differentiation was induced by changing to differentiation medium containing PBS or IFN-γ (25 ng/ml). The medium with PBS or IFN-γ was refreshed daily.

To determine its effect *in vivo*, WT C57BL/6N mice were injected intramuscularly with 7500 U of Recombinant Mouse IFN-γ (R&D Systems #485-MI). The contralateral leg was injected with PBS as vehicle control.

### In vivo IFN-γ neutralization

MCM;FLExD mice were injected with TMX (final concentration of 5 mg/kg) at d0 and d1. Additionally, mice were IP-injected with Purified in vivo GOLD Functional Grade Armenian Hamster Anti-Mouse IFN-γ (Leinco Technologies Clone H22 # I-438) or Purified in vivo GOLD Functional Grade Armenian Hamster IgG Isotype Control (Leinco Technologies Clone PIP #I-140), 200μg dissolved in 100μl of PBS, every three days. Mice were sacrificed 15 days post DUX4 induction.

### siRNA transfection

*DUX4* and non-silencing control siRNAs (listed in Table S8) were purchased from Dharmacon and used at a final concentration of 25 nM. Transfection of siRNAs was performed by using Lipofectamine 3000 Transfection Reagent (Invitrogen #L3000-008) following the manufacturer’s instructions.

### Luminex Assay

Luminex was performed as in (35). Briefly, muscle homogenates were prepared from freshly dissected Gastrocnemius muscle with the sample-grinding kit (Sigma-Aldrich #GE80-6483-37). Muscle homogenates were normalized to a concentration of 2000 μg/ml. 50μl of sample (1:2 diluted) were used and mixed to the microparticle cocktail. The Luminex Discovery Assay was performed following the instruction provided by Bio-Techne (Mouse Premixed Multi-Analyte Kit #LXSAMSM-05). Fluorescence was quantified using a Luminex 200 instrument.

### Nuclei isolation and snRNA sequencing

From each mouse, nuclei were isolated from a pool of two freshly dissected Tibialis Anterior (TA) muscles, which were kept in ice-cold PBS + RNase inhibitor (0.2 U/μl) (Sigma-Aldrich #3335399001) in a 2 ml tube. Every step of single nuclei isolation was performed on ice or at 4°C. Nuclei were isolated following the instructions from (139). Briefly, after muscles dissection, the tissue was cut for 2 min in ice-cold lysis buffer and then lysed for additional 3 min on ice with gentle shaking. Then, the preparation was transferred in a Dounce homogenizer on ice and gently dounced for 10 times. Nuclei were then filtered sequentially with 70 μm, 40 μm, 30 μm cell strainers and centrifuged for 5 min at 500g at 4°C. After wash in wash buffer + RNase inhibitor (0.2 U/μl), to the resuspended pellet was added DAPI (Sigma-Aldrich #D9542) (10 μg/ml) for 5 min in the dark for nuclei visualization by FACS. FACS sorting of the nuclei was performed as in (139). The quality and concentration of sorted nuclei were verified on a fluorescent microscope. Single nuclei encapsulation, cDNA amplification, library prep and sequencing were performed by the San Raffaele Center for Omic Sciences (COSR, Milan-Italy). 10.000 nuclei were loaded onto a single-cell chip for GEM generation using the 10x Genomics Chromium Controller (10× Genomics, Pleasanton, CA). 3′mRNA-seq gene expression libraries were constructed using the Chromium Single Cell 3′ Library & Gel Bead Kit v3.1 (10x Genomics) according to the manufacturer’s guidelines. Briefly, nuclei suspensions were diluted in PBS with 0.04% BSA to a final concentration of 1 × 10^6^ nuclei/ml (1.000 nuclei per μl). All reactions were performed in a C1000 Touch Thermal Cycler (Bio-Rad Laboratories, Hercules, CA). Twelve cycles were used for cDNA amplification, while the number of total cycles for the sample index PCR was calculated based on the cDNA concentration. Amplified cDNA and final libraries were evaluated using a Tape Station 4200 (Agilent Technologies, Santa Clara, CA) with a high sensitivity chip. snRNA-seq libraries were sequenced on an Illumina NovaSeq6000 with the standard sequencing protocol. snRNA-seq was performed on one ACTA1-MCM and three MCM;FLExD mice at 15 days post TMX induction.

### snRNA-seq analysis

Raw feature-barcode matrices were processed with CellBender v0.3.2 to remove ambient RNA contamination and empty droplets (140). Quality control was performed in R using the scCustomize and Seurat v5 frameworks (141,142), retaining nuclei with <10% mitochondrial content and >200 detected genes, with sample-specific upper gene-count thresholds applied to exclude potential multiplets. Data were log-normalized and the top 2,000 highly variable genes were selected using the variance-stabilizing transformation method. Batch effects were corrected using the Harmony algorithm (143). Initial cell-type identities were assigned by marker gene identification using the Wilcoxon rank-sum test and validated through muscle dataset reference inspection (144–150). Potential doublets were identified and removed using scDblFinder (151), after which the dataset was re-processed from scratch to ensure a clean low-dimensional embedding. Fibro-adipogenic progenitors (FAPs) and macrophages were further sub-clustered to identify high-resolution cell states. After sub-clustering, contaminant clusters of myofibers were removed from both FAPs and macrophages analysis. Muscle fiber nuclei were similarly subset for refined analysis. Transcriptional activity was quantified using gene module scoring. A curated 17-gene DUX4 target signature (Table S6) and the Mouse Gene Set: HALLMARK_INTERFERON_GAMMA_RESPONSE (152) were scored across all nuclei using the AddModuleScore function (153). Violin with represents density; increased smoothing (bandwidth adjustment = 3 or 4) was applied to reduce artifacts from discrete scoring at lower values.

Statistical comparisons between fiber types and experimental conditions were performed using pairwise Wilcoxon rank-sum tests. All visualizations, including UMAPs, heatmaps, and violin plots, were generated using Seurat, pheatmap, and ggplot2 (154).

### Spatial transcriptomics

TA muscles from ACTA1-MCM and MCM;FLExD mice at 2-6-15 days post TMX induction were immediately dissected and frozen in optimal cutting temperature (OCT) (Sakura #4583) embedding media in liquid nitrogen–cooled isopentane. Muscle tissue sections were cryosectioned at -20°C at 10 μm thickness onto Superfrost PLUS Adhesion Microscope slides (Epredia #J1800AMNZ), following the protocol 10X Genomics CG000636 | RevA and stored at −80°C for one night. The following day the samples were delivered and processed for the next steps by COSR (Milan-Italy). Samples were processed as described in 10x Genomics CG000685_VisiumHD_GeneExpression_UserGuide_RevD and CG000763_VisiumHDFFTissuePrepHandbook_RevA. The 10x Genomics Visium HD Spatial Gene Expression platform was used for spatial transcriptomics analysis according to the manufacturer’s guidelines.

### Spatial transcriptomics analysis

Tissue sections were processed according to the manufacturer’s instructions and aligned to the GRCm38/mm10 mouse reference genome using Space Ranger vX (10x Genomics, 2024). All downstream analyses were performed at 8 µm bin resolution to maximize spatial precision. To manage the high-resolution data, raw gene-expression matrices were stored as on-disk BPCells directories (155). Quality control was performed on the merged dataset, where bins with fewer than 90 total UMI counts were removed to filter background signal. Data were library-size normalized using the median total UMI count across all bins as a scale factor, followed by log1p transformation. The top 2,000 highly variable genes were identified using the ‘vst’ selection method (141). To ensure computational scalability, a sketch-based subsampling strategy was employed where approximately one-quarter of the bins per sample were selected using leverage-score-based sketching (141). Dimensionality reduction was performed on this sketched subset using PCA, retaining the first 30 principal components for downstream steps. Batch correction was applied using the Harmony algorithm (143). Clustering was performed on the Harmony-corrected embedding using a shared nearest-neighbor graph and the Louvain algorithm at a resolution of 0.5. The resulting cluster labels and UMAP coordinates were then projected back to the full dataset using the ProjectIntegration and ProjectData frameworks (141). Cluster marker genes were identified using the Wilcoxon rank-sum test, and cell-type identities were assigned through a combination of automated LLM-assisted proposals and manual curation against established skeletal muscle atlases, yielding 14 distinct cell types. To mitigate integration-driven artifacts, a spatial connectivity quality-control step was implemented. Following the integration of the Visium HD spatial transcriptomics samples into a single merged object, we performed a spatial connectivity analysis to identify and exclude any spurious cell type annotations introduced by the harmonization process. Specifically, since the DUX4 transgene is absent in ACTA1-MCM mice, any pixel annotated as a “DUX4-induced aberrant myofiber” in that condition is expected to represent a misclassification artifact rather than a true biological signal. To systematically identify such noise pixels across all samples, we extracted spatial coordinates for each pixel from the individual pre-integration Seurat objects and computed pairwise Euclidean distances between all pixels assigned to the DUX4-induced aberrant myofiber identity. The grid spacing of each sample was estimated from a random subset of pixels, and the number of immediately adjacent neighbors sharing the same cell type identity (within 1.5× the estimated grid spacing) was computed for each DUX4-induced aberrant myofiber-annotated pixel. The distribution of DUX4-induced aberrant myofiber neighbor counts was then inspected for each sample independently. Since the ACTA1-MCM sample cannot contain true DUX4-induced aberrant myofibers by design, the full range of neighbor counts observed in that sample (0–19) was used as an empirical, data-driven threshold to define noise pixels: any pixel across all four samples with 19 or fewer DUX4-induced aberrant myofiber neighbors was classified as isolated and likely artifactual. To further validate this threshold, we tested the Spearman correlation between spatial connectivity and the DUX4 transcriptional target module score for each sample independently. Consistent with the noise interpretation, no significant correlation was observed in the control sample, indicating that DUX4-annotated pixels in that condition are spatially and transcriptionally unstructured. In contrast, all MCM;FLExD samples showed highly significant positive correlations, indicating spatial coherence consistently and significantly associated with DUX4 transcriptional activity. This threshold was empirically defined by the maximum neighbor count observed in the uninduced normal sample. Transcriptional signature enrichment was quantified using the UCell R package, which utilizes a rank-based Mann-Whitney U statistic (156). To generate the Violin plot showing the UCell Signature scores, only non-zero scores are visualized to emphasize the distribution of pixels with detectable DUX4 target gene expression of IFN-γ pathway activity; zeros (pixels with no expression of signature genes) were retained in statistical comparisons. Scores were computed for the 17-gene DUX4 target signature set, a 33-gene Type 2B myofiber signature derived from cluster marker genes, and the Mouse Gene Set: HALLMARK_INTERFERON_GAMMA_RESPONSE (152). Following initial cell-type assignment, we observed that Harmony-based joint clustering, while correctly identifying the major transcriptional states, may conservatively retain in the Type 2B cluster a subset of bins from DUX4-expressing samples whose global transcriptome still partially resembles healthy fast-twitch fibers, yet whose signature scores indicate incipient transcriptional reprogramming. To address this, a post-hoc score-based reclassification was applied exclusively to bins initially assigned as Type 2B. Bins were reclassified as DUX4-induced aberrant myofibers if their DUX4 target UCell score met or exceeded an empirically determined threshold (≥ 0.1), chosen to be intentionally non-stringent so as to capture pixels with measurable DUX4 target activation while minimizing false reclassification of truly healthy fibers. Threshold selection was guided by visual inspection of per-sample score distributions, including marginal histograms and joint scatter plots of the DUX4 target score versus the Type 2B signature score within Type 2B-assigned bins. As a key sanity check, the proportion of reclassified bins in the uninduced ACTA1-MCM control was verified to be negligible, confirming that the threshold does not introduce spurious reclassification in the absence of DUX4 induction. The revised cell-type labels were stored as a separate metadata field (cell_types_full2) and used for all subsequent compositional and comparative analyses. Pairwise statistical comparisons between groups were performed using the Wilcoxon rank-sum test with Cohen’s d used as an effect-size measure. Final visualizations including spatial tile maps and feature plots were generated using custom ggplot2 frameworks (154) and 10x Genomics Loupe Browser v8.

### RNA extraction and RT-qPCR

Total RNA from isolated muscles was extracted with TRIzol Reagent (Invitrogen #15596-018) using manufacturer’s instructions. Total RNA from human cells was extracted using PureLink RNA Mini Kit (Invitrogen #12183025) following manufacturer’s instructions. cDNA synthesis was performed using Super Script III First-Strand Synthesis Super-Mix (Thermo Fisher Scientific #11752-250) following manufacturer’s instructions. Quantitative real-time PCR (qPCR) was performed with iTaq Universal SYBR Green Supermix (Biorad #1725121) using CFX96 or CFX384 Real-Time PCR Detection Systems (Bio-Rad). Primers used for RT-qPCR are listed in Table S8. Relative quantification was performed using the ΔΔCt method. *GAPDH* was used as housekeeping gene for sample normalization. For each group of cells (human primary muscle cells from healthy or FSHD patient donors), the fold change was quantified based on the average of the 2^-ΔCt of the healthy donor group. For mouse samples, the fold change was quantified based on the average of the 2^-ΔCt of the ACTA1-MCM group. Specific details of each data set are provided in the Figure legends.

### Analysis of publicly available gene expression datasets

We analyzed gene expression profiling from six independent human cohorts for a total of 47 healthy donors (59 muscle biopsies) and 154 FSHD patients (218 muscle biopsies). Banerji *et al*., EGAS00001007350 (26), describe 59 muscle biopsies assessed by RNA-seq: 11 from control individuals and 48 from FSHD patients. The FSHD patients underwent two muscle biopsies each, one from a non-inflamed muscle (TIRM− on MRI) and one from an inflamed muscle (TIRM+ on MRI), resulting in 24 TIRM− samples and 24 corresponding isogenic TIRM+ samples. Wang *et al*., GSE115650 (28), describe 43 muscle biopsies assessed by RNA-sequencing: nine from control individuals and 34 from FSHD patients. For the FSHD patients, 23 biopsies are from a non-inflamed muscle, and 11 biopsies are from an inflamed muscle based on histological analyses. Wong *et al*., GSE140261 (29), describe 35 muscle biopsies assessed by RNA-seq: 8 from control individuals and 27 from FSHD patients. For the FSHD patients, 17 biopsies are from a non-inflamed muscle, and 10 biopsies are from an inflamed muscle based on histological analyses. van den Heuvel *et al.,* EGAS00001005782 (31), describes 61 muscle biopsies assessed by RNA-seq: 28 FSHD patients (37 biopsies) and 12 control individuals (24 biopsies). Wong et al., GSE242912 (23), describe 64 muscle biopsies assessed by RNA-seq: 33 FSHD patients, with bilateral muscle biopsies from 31 patients. 56 biopsies are from a non-inflamed muscle, and 8 biopsies are from an inflamed muscle based on histological analyses. Tasca *et al*., GSE26852 (27), describe 15 muscle biopsies assessed using microarray: 7 from control individuals and 8 from FSHD patients. For the FSHD patients, 4 biopsies are from a non-inflamed muscle (T2-STIR− on MRI) and 4 biopsies are from an inflamed muscle (T2-STIR+ on MRI).

Jones *et al*., GSE122562 (30), describe 12 muscle biopsies assessed by RNA-seq: 3 from ACTA1-MCM mice, 3 from MCM;FLExD with a mild phenotype, 3 from MCM;FLExD with a moderate phenotype, and 3 from MCM;FLExD with a severe phenotype.

For all datasets except EGAS00001005782, EGAS00001007350, GSE242912 and GSE122562, differentially expressed genes (DEGs) where identified using the NCBI GEO2R web tool (https://www.ncbi.nlm.nih.gov/geo/info/geo2r.html) using default parameters.

For EGAS00001005782, raw gene-level count data and associated sample metadata were kindly provided by Drs. Anita van den Heuvel and Silvère M. van der Maarel. For EGAS00001007350, upregulated genes were obtained directly from Table S1 of (56). For GSE242912, gene expression count matrices were retrieved from NCBI GEO (https://www.ncbi.nlm.nih.gov/geo/). Differential gene expression analysis was conducted between inflamed FSHD muscles compared to not inflamed FSHD muscles samples, using the edgeR package v4.6.3 (157). For GSE122562, upregulated genes were obtained directly from Table S2 of (30).

The list of significantly (adjusted p-value ≤ 0.05) upregulated genes in the various comparisons for the different datasets is present in Table S1.

For all datasets, to perform Molecular Signatures Database (MSigDB) (158,159) Hallmark gene sets pathways enrichment analysis, significantly upregulated genes were analyzed with the Enrichr web tool (https://maayanlab.cloud/Enrichr/) (160–162).

Significantly (adjusted p-value ≤ 0.05) enriched pathways are reported in Table S2.

To perform Gene Set Enrichment Analyses (GSEA), RNA-seq count matrices from individual studies (GSE115650, GSE122562, GSE140261, GSE242912) were retrieved from GEO. For GSE115650 and GSE242912, samples were categorized as inflamed (score > 0) or not inflamed (score = 0) based on the reported inflammation score in the metadata. Samples with missing inflammation score values were removed from downstream analyses. Differential gene expression analysis was separately conducted between FSHD vs. healthy donor human samples, inflamed FSHD vs. not inflamed FSHD human samples, or FLExDUX4 vs. ACTA1-MCM mouse samples using the edgeR package v4.6.3 (157). For all the datasets, samples with significant deviation from clustering in the PCA were identified as outliers and excluded from the analysis. GSEA was performed using normalized expression values obtained from edgeR, alongside species-appropriate gene sets retrieved from the MSigDB (158,159), via the GSEA tool (https://www.gsea-msigdb.org/gsea/index.jsp) v4.4.0 (163). The phenotype file was used to assign the corresponding labels to the samples in the expression table, and Ensembl IDs were collapsed to gene symbols using human/mouse Ensembl_Gene_ID_MSigDB v2026 chip. The GSEA results are reported in Table S3.

### DUX4 signature target genes in mouse skeletal muscle

To generate the list of putative direct DUX4 target genes in mouse muscle, we matched RNA-seq and ChIP-seq datasets for DUX4 in mouse muscle cells [(105); GSE87282] with the list of significantly upregulated genes in moderate and severe MCM;FLExD compared to ACTA1-MCM control mice [(30); GSE122562]. This produced a list of putative direct DUX4 target genes in mouse skeletal muscle, and the 17 top expressed genes were subsequently used as DUX4 target signature set (Table S6) to identify muscle fibers with active DUX4 in snRNA-seq and ST.

### FACS analysis

Cell isolation and labeling from muscle tissue were performed essentially as described in (164). Briefly, the single cell suspension was stained with Live/Dead fixable Aqua Dead cell stain kit (Invitrogen #L34957) 1:100 in PBS for 10 min at 4°C to identify live cells. Then, after one wash in HBSS (Gibco #14025100) + 0.2% BSA (Sigma-Aldrich #A3912) + 1% P/S, the single cell suspension was stained with CD31 (Miltenyi Biotec #130-111-539)/CD45(Miltenyi Biotec #130-110-795)/TER119 (Miltenyi Biotec #130-112-907) Biotin (1:25) and Anti-Biotin-VioBlue (Miltenyi Biotec #130-110-958 1:50) for lineage exclusion for 40 min at 4°C. From Lineage negative, FAPs were defined as Sca1-FITC positive cells (Miltenyi Biotec #130-116-490; 1:25) and α7-integrin-APC negative cells (Miltenyi Biotec #130-123-833; 1:25).

For Desmin staining of human muscle cells, cells were harvested and washed to obtain a single-cell suspension. Cell concentration was adjusted to 1–5 × 10⁶ cells/mL in 500 µL of ice-cold MACS buffer (PBS supplemented with 0.5% BSA and 2 mM EDTA). Cells were incubated in blocking buffer (MACS buffer supplemented with 2% BSA and 5% horse serum) for 30 min on ice. Cells were then incubated with primary antibody against Desmin (Leica #DES-DERII-L-CE, 1:50) for 30 min at RT. Following incubation, cells were washed three times with MACS buffer and resuspended in 200 µL of ice-cold MACS buffer. Subsequently, cells were incubated with the appropriate fluorochrome-conjugated secondary antibody (Goat anti-Mouse IgG Alexa Fluor 488, Invitrogen #A11029) diluted in MACS buffer for 30 min at RT. Cells were washed three times with MACS buffer and finally resuspended in 200 µL of ice-cold MACS buffer. Cells were sorted using the BD FACSAria Fusion (BD) and analyzed at BD FACSCANTO II (Becton Dickinson). Data were analyzed using FlowJo v10.7 (BD Life Sciences).

### Histological analysis

For hematoxylin and eosin staining, frozen sections were fixed in 4% PFA for 30 min, at room temperature (RT). Then, the staining was performed as per the 10X Genomics protocol (CG000763 RevA from Visium HD Spatial Gene Expression Reagent kits). Images were collected using an AxioImage M2 fluorescent microscope (Zeiss, Germany).

### Immunofluorescence

iDUX4 and iVector cells grown on glass coverslips were fixed with 4% PFA in PBS for 10 min at RT. Cells were then permeabilized with 0.5% Triton X-100 in PBS for 20 min at RT. To reduce non-specific binding, samples were blocked with 5% BSA (Sigma-Aldrich #A3912) in PBS for 1h at RT. Primary antibodies incubation was performed overnight at 4 °C using the following antibodies diluted 1:100 in blocking buffer: mouse anti-double-stranded RNA antibody K1 (Nordic-MUBio #10020500) and rabbit anti-DUX4 (Abcam #ab124699). After three washes for 10 min in PBS 1X, the cells were incubated for 45 min at RT with the appropriate secondary antibodies diluted 1:500 in blocking solution: Goat anti-mouse IgG Alexa Fluor 488 (Invitrogen #A32723) and Goat anti-rabbit IgG Alexa Fluor 555 (Invitrogen #A21429). Nuclei were counterstained with DAPI (Sigma-Aldrich #D9542) diluted 1:1000 in PBS. Coverslips were mounted using FluorSave Reagent (Millipore #345789).

For myotube morphology analysis, differentiated primary healthy donor muscle cells treated with vehicle or recombinant IFN-γ were fixed in 4% PFA for 10 min at RT. Cells were incubated with blocking buffer containing 5% Goat serum and 0.1% Triton X-100 for 1h at RT. Then, cells were incubated with primary antibody anti-myosin heavy chain (Developmental Studies Hybridoma Bank, MF-20) for 45 min at RT, and then incubated with Goat anti-mouse IgG Alexa Fluor 488 (Invitrogen #A32723) diluted 1:500 and DAPI (Sigma-Aldrich #D9542) in blocking buffer for 45 min at RT. The differentiation index was scored by determining the percentage of myosin heavy chain-positive (MHC+) nuclei. For each treatment, three differentiation experiments were performed and at least 5 fields per well were analyzed.

For immunofluorescence on tissue sections, muscles were snap-frozen in liquid nitrogen–cooled isopentane (Sigma-Aldrich, #M32631) and stored at -80°C until sectioning. 10 μm transverse cryosections were cut from Tibialis Anterior (TA) or Gastrocnemius (GA). All the tissue sections were fixed in 4% PFA for 10 min. For Laminin, Col1a and Perilipin and Necrotic staining, tissue sections were permeabilized with 100% methanol for 6 min at −20°C and blocked with 5% BSA in PBS for 1 hour at RT to avoid nonspecific binding. For CD68 and PDGFR*α* staining, cryosections were permeabilized with 0.2% Tryton X-100 in PBS for 20 min at RT. Immunostaining with primary antibodies was performed overnight at 4°C. CD68 (MCA #1957GA), Laminin (Sigma-Aldrich #L9393), Col1a (Santa Cruz #sc59772), Perilipin (Sigma-Aldrich #P1873) diluted 1:300 in blocking buffer, PDGFRα (R&D Systems #AF1062), F4/80 (Abcam #300421) diluted 1:100 in blocking buffer. Then, sections were incubated for 45 min at RT with the appropriate secondary antibodies: Goat anti-mouse IgG Alexa Fluor 555 (Invitrogen #A21424), Goat anti-rabbit IgG Alexa Fluor 488 (Invitrogen #A11034), Donkey anti-goat Alexa Fluor 594 (Invitrogen #A11058) 1:500 in blocking buffer. Necrotic myofibers were stained using Goat anti-mouse IgG Alexa Fluor 488 (Invitrogen #A32723). Nuclei were counterstained with DAPI (Sigma-Aldrich, #D9542) 1:1000 in PBS. For dsRNA staining, following fixation, slides were washed and then permeabilized in 0.1% Triton X-100 in PBS for 20 min at RT. Blocking was performed using Mouse Ig Blocking Reagent (from M.O.M. Immunodetection Kit, Vector Labs #BMK-2202), diluted in PBS according to the manufacturer’s instructions, for 1h at RT. After blocking, slides were washed and incubated for 5 min in M.O.M. Diluent. Primary antibodies were incubated overnight at 4°C, diluted 1:100 in M.O.M. diluent: mouse anti-dsRNA K1 (Nordic-MUBio #10020200). Then sections were incubated for 45 min at RT with the appropriate secondary antibodies: Goat anti-mouse IgG Alexa Fluor 555 (Invitrogen #A21424) diluted 1:500 in M.O.M. diluent. Nuclei were counterstained with DAPI (Sigma-Aldrich #D9542) diluted 1:1000 in PBS. Sections were mounted with FluorSave Reagent (Millipore #345789). Images were collected using an AxioImage M2 fluorescent microscope (Zeiss, Germany).

### RNase treatment for dsRNA staining specificity

To assess the specificity of the dsRNA immunofluorescence signal, treatments with RNases degrading double-stranded RNA or single-stranded RNA, as negative control, were performed prior to the blocking step using a protocol adapted from (165). Briefly, for double-stranded RNA degradation samples were treated with a reaction mix containing 2 µL ShortCut RNase III (New England Biolabs #M0245S), 5 µL reaction buffer, 5 µL MgCl₂, and 38 µL nuclease-free H₂O. For single-stranded RNA degradation, samples were treated with a reaction mix composed of 2 µL RNase A/T1 (Thermo Fisher Scientific #EN0551), 5 µL reaction buffer, and 43 µL nuclease-free H₂O. For each treatment, 50 µL of the corresponding reaction mix were applied to the samples, which were then incubated for 30 minutes at 37°C. Following incubation, samples were washed in PBS and processed for immunofluorescence staining as above.

### RNAscope combined with immunofluorescence

To combine RNAscope in situ hybridization with immunofluorescence, the RNAscope Multiplex Fluorescent v2 Assay combined with immunofluorescence protocol (Advanced Cell Diagnostics, ACD 323100-TN/RevA) was followed. RNAscope in situ hybridization was performed using the RNAscope multiplex Fluorescent Reagent Kit v2 Assay following the manufacturer’s instructions (RNAscope Multiplex Fluorescent Reagent Kit v2 User Manual, Advanced Cell Diagnostics, ACD #323100-USM). Briefly, the fixed-frozen tissue sample preparation and pretreatment protocols were followed for muscle tissue. Gastrocnemius muscles were dissected and fixed in freshly prepared 4% PFA for 24h at 4°C. Next, the muscles were immersed in a gradient scale of sucrose 10-20-30% in PBS at 4°C until the tissue sunk to the bottom of the container. Then, muscles were frozen in Optimal Cutting Temperature (OCT) embedding media with liquid nitrogen–cooled isopentane and stored at -80 C° until sectioning. Serial 5 μm transverse cryosections were cut and, upon tissue dehydration, the slides were immersed in the target retrieval 1X buffer for 5 min at 99°C. Then, the slides were air dried briefly and boundaries were drawn around each section using a hydrophobic pen. When hydrophobic boundaries had dried, the Protease III reagent was added to each section until fully covered, and the slides were incubated 30 minutes at 40°C. Each slide was then placed in a prewarmed humidity control tray (ACD) containing dampened filter paper and mouse *IFN-*γ probes (mmIFN-γ ACD #311391-C2) were added onto each section until fully submerged. The humidity control tray was placed in a HybEZ oven (ACD) for 2 hours at 40°C. Following incubation with the probe, the slides were washed two times in RNAscope wash buffer and returned to the oven for 30 minutes after submersion in AMP-1 reagent. Washes and amplification were repeated using AMP-2 and AMP-3 reagents with a 30 min or 15 min incubation period, respectively. Signal revelation was performed with the TSA Vivid Fluorophore Kit 650 (ACD #323273). The slides were then processed for immunofluorescence following the protocols suggested from ACD. The slides were stained with anti-Laminin (1:300) (Sigma-Aldrich #L9393) for 2h at RT and, after washes, the slides were stained with Goat anti-rabbit IgG Alexa Fluor 488 (Invitrogen #A11034) 1:500 in TBS 0,1% BSA for 30 min at RT and then incubated in DAPI (Sigma-Aldrich #D9542) diluted 1:1000 in PBS for 1 min, and cover-slipped with Prolong Gold Antifade mounting medium (Invitrogen #P36930). Images were collected using a Leica Confocal SP5 microscope (Leica Microsystems). The quantification was performed with the QuPath software (166) by the quantification of the single dots per area following ACD Tech Note MK-51-154.

### Use of AI-Assisted Technologies

For the analysis of snRNA-Seq and ST results, Claude (version 3.5 Sonnet) was used for assistance in generating R code for data visualization and Gemini Flash (Google) was used for aid in biological marker interpretation. These tools were used to support the analytical workflow and all AI-suggestions were manually verified, edited, and validated by the authors against established literature and biological databases. The authors maintain full responsibility for the scientific integrity of the work.

### Quantification and statistical analyses

For image quantification, analyses were performed on at least five distinct fields per sample to ensure representative coverage of the entire transverse muscle section. The dsRNA positive area, Perilipin-positive droplets and Collagen infiltration were quantified using ImageJ, calculating the area of the pixels (square pixel) per field. The muscle fiber cross-sectional area (CSA) was calculated using an ImageJ software plug-in (Macro_seg_5_modif.ijm.txt). Areas were measured on images taken randomly to cover the entire transverse area of the muscle section. The necrotic myofibers were counted over the total number of the myofibers in the field. The CD68, PDGFRα and F4/80 staining quantification was performed as the percentage of positive cells over the total number of cells. For each field, positive cells were manually counted on Fiji, while total nuclei were automatically assessed via the count_nuclei.ijm macro of ImageJ.

All statistical analyses were performed using GraphPad Prism 9 (GraphPad Software, San Diego, USA). Quantitative data are presented as mean ± standard deviation (SD). Statistical significance was calculated on at least three independent experiments. P-value: *p < 0.05; **p < 0.01; ***p < 0.001; ****p < 0.0001. The details of the statistical test used for each dataset are provided in the corresponding figure legend.

## Acknowledgments

We would like to thank Silvère M. van der Maarel and Anita van den Heuvel for the expression counts table and metadata for EGAS00001005782. We are grateful to Baziel van Engelen, Peter Zammit and Anna Greco for granting access to EGAS00001007350. We thank Rabi Tawil and the FSHD Research Center at UR Medicine, and Charles P Emerson and the UMass Chan Wellstone Center for FSHD for primary and immortalized muscle cells from healthy donors and FSHD patients. We are grateful to Peter L Jones and Andreia M Nunes for the conditions to perform muscle extracts for Luminex assays. We thank Silvia Gregori and Giada Amodio for the help to perform the Luminex assay. We are grateful to Monica Nicolau, Chiara Nicoletti and Pier Lorenzo Puri for the helpful discussion about snRNA-seq analysis. We thank Anna Kajaste-Rudnitski and Giovanni Tonon for critically reading the manuscript.

## Author contributions

Conceptualization: DG, BB Methodology: BB, DG, JMGM

Investigation: BB, FDP, MS, JMGM, RA, EF, AV, BC, FT, İE, DB, MM, DC, MP, EM, GS, SE, GR

Visualization: BB, JMGM

Funding acquisition: DG, BB, GS, SE Project administration: DG, BB Supervision: DG

Writing – original draft: BB, DG Writing – review & editing: BB, DG

## Data, code, and materials availability

All data are available in the main text or the supplementary materials.

snRNA-seq and ST data have been deposited in the GEO database with accession number GSE327524.

All reasonable requests for materials will be fulfilled upon completion of a material transfer agreement.

## REFERENCES

1. Mocciaro E, Runfola V, Ghezzi P, Pannese M, Gabellini D. DUX4 Role in Normal Physiology and in FSHD Muscular Dystrophy. Cells. 2021 Dec 1;10(12). doi:10.3390/CELLS10123322 PubMed PMID: 34943834.

2. Cohen J, DeSimone A, Lek M, Lek A. Therapeutic Approaches in Facioscapulohumeral Muscular Dystrophy. Trends Mol Med. 2021 Feb 1;27(2):123–37. doi:10.1016/J.MOLMED.2020.09.008 PubMed PMID: 33092966.

3. Feng Q, Snider L, Jagannathan S, Tawil R, van der Maarel SM, Tapscott SJ, et al. A feedback loop between nonsense-mediated decay and the retrogene DUX4 in facioscapulohumeral muscular dystrophy. Elife. 2015 Jan 7;4(4). doi:10.7554/ELIFE.04996 PubMed PMID: 25564732.

4. Campbell AE, Shadle SC, Jagannathan S, Lim JW, Resnick R, Tawil R, et al. NuRD and CAF-1-mediated silencing of the D4Z4 array is modulated by DUX4-induced MBD3L proteins. Elife. 2018 Mar 13;7. doi:10.7554/ELIFE.31023 PubMed PMID: 29533181.

5. Resnick R, Wong CJ, Hamm DC, Bennett SR, Skene PJ, Hake SB, et al. DUX4-Induced Histone Variants H3.X and H3.Y Mark DUX4 Target Genes for Expression. Cell Rep. 2019 Nov 12;29(7):1812–1820.e5. doi:10.1016/J.CELREP.2019.10.025 PubMed PMID: 31722199.

6. Jiang S, Williams K, Kong X, Zeng W, Nguyen NV, Ma X, et al. Single-nucleus RNA-seq identifies divergent populations of FSHD2 myotube nuclei. PLoS Genet. 2020 May 1;16(5):e1008754. doi:10.1371/JOURNAL.PGEN.1008754 PubMed PMID: 32365093.

7. Chau J, Kong X, Viet Nguyen N, Williams K, Ball M, Tawil R, et al. Relationship of DUX4 and target gene expression in FSHD myocytes. Hum Mutat. 2021 Apr 1;42(4):421. doi:10.1002/HUMU.24171 PubMed PMID: 33502067.

8. Hamm DC, Paatela EM, Bennett SR, Wong CJ, Campbell AE, Wladyka CL, et al. The transcription factor DUX4 orchestrates translational reprogramming by broadly suppressing translation efficiency and promoting expression of DUX4-induced mRNAs. PLoS Biol. 2023 Sep 1;21(9). doi:10.1371/JOURNAL.PBIO.3002317 PubMed PMID: 37747887.

9. Arends T, Tsuchida H, Adeyemi RO, Tapscott SJ. DUX4-induced HSATII transcription causes KDM2A/B-PRC1 nuclear foci and impairs DNA damage response. J Cell Biol. 2024 May 6;223(5). doi:10.1083/JCB.202303141 PubMed PMID: 38451221.

10. Arends T, Bennett SR, Tapscott SJ. DUX4-induced HSATII RNA accumulation drives protein aggregation impacting RNA processing pathways. J Cell Biol. 2025 Feb 2;225(2):e202501129. doi:10.1083/JCB.202501129 PubMed PMID: 41329166.

11. Brennan CM, Hill AS, St. Andre M, Li X, Madeti V, Breitkopf S, et al. DUX4 expression activates JNK and p38 MAP kinases in myoblasts. Dis Model Mech. 2022 Nov 1;15(11). doi:10.1242/DMM.049516 PubMed PMID: 36196640.

12. Dahlqvist JR, Andersen G, Khawajazada T, Vissing C, Thomsen C, Vissing J. Relationship between muscle inflammation and fat replacement assessed by MRI in facioscapulohumeral muscular dystrophy. J Neurol. 2019 May 1;266(5):1127–35. doi:10.1007/S00415-019-09242-Y PubMed PMID: 30778707.

13. Tidball JG, Welc SS, Wehling-Henricks M. Immunobiology of Inherited Muscular Dystrophies. Compr Physiol. 2018 Sep 14;8(4):1313–56. doi:10.1002/CPHY.C170052 PubMed PMID: 30215857.

14. Frisullo G, Frusciante R, Nociti V, Tasca G, Renna R, Iorio R, et al. CD8+ T cells in facioscapulohumeral muscular dystrophy patients with inflammatory features at muscle MRI. J Clin Immunol. 2011;31(2). doi:10.1007/s10875-010-9474-6

15. Janssen BH, Voet NBM, Nabuurs CI, Kan HE, De Rooy JWJ, Geurts AC, et al. Distinct disease phases in muscles of facioscapulohumeral dystrophy patients identified by MR detected fat infiltration. PLoS One. 2014 Jan 14;9(1). doi:10.1371/JOURNAL.PONE.0085416 PubMed PMID: 24454861.

16. Dahlqvist JR, Poulsen NS, Østergaard ST, Fornander F, De Stricker Borch J, Danielsen ER, et al. Evaluation of inflammatory lesions over 2 years in facioscapulohumeral muscular dystrophy. Neurology. 2020 Sep 1;95(9):E1211–21. doi:10.1212/WNL.0000000000010155 PubMed PMID: 32611642.

17. Tawil R, McDermott MP, Pandya S, King W, Kissel J, Mendell JR, et al. A pilot trial of prednisone in facioscapulohumeral muscular dystrophy. FSH-DY Group. Neurology. 1997 Jan;48(1):46–9. PubMed PMID: 9008492.

18. Kissel JT, McDermott MP, Mendell JR, King WM, Pandya S, Griggs RC, et al. Randomized, double-blind, placebo-controlled trial of albuterol in facioscapulohumeral dystrophy. Neurology. 2001 Oct 23;57(8):1434–40. PubMed PMID: 11673585.

19. Van Der Kooi EL, Kalkman JS, Lindeman E, Hendriks JCM, Van Engelen BGM, Bleijenberg G, et al. Effects of training and albuterol on pain and fatigue in facioscapulohumeral muscular dystrophy. J Neurol. 2007 Jul;254(7):931–40. doi:10.1007/S00415-006-0432-4 PubMed PMID: 17361345.

20. Payan CA, Hogrel JY, Hammouda EH, Lacomblez L, Ollivier G, Doppler V, et al. Periodic salbutamol in facioscapulohumeral muscular dystrophy: a randomized controlled trial. Arch Phys Med Rehabil. 2009 Jul;90(7):1094–101. doi:10.1016/J.APMR.2008.12.027 PubMed PMID: 19577021.

21. Kissel JT, McDermott MP, Natarajan R, Mendell JR, Pandya S, King WM, et al. Pilot trial of albuterol in facioscapulohumeral muscular dystrophy. Neurology. 1998;50(5):1402–6. doi:10.1212/WNL.50.5.1402

22. Bloch RJ, Traficante M, Mariot V, Dumonceaux J. The recent clinical trial of losmapimod for the treatment of facioscapulohumeral muscular dystrophy. Neuromuscul Disord. 2025 Jul 1;52. doi:10.1016/J.NMD.2025.105422 PubMed PMID: 40580827.

23. Wong CJ, Friedman SD, Snider L, Bennett SR, Jones TI, Jones PL, et al. Regional and bilateral MRI and gene signatures in facioscapulohumeral dystrophy: implications for clinical trial design and mechanisms of disease progression. Hum Mol Genet. 2024 Apr 15;33(8):698–708. doi:10.1093/HMG/DDAE007 PubMed PMID: 38268317.

24. Greco A, Straasheijm KR, Mul K, Van Den Heuvel A, Van Der Maarel SM, Joosten LAB, et al. Profiling Serum Antibodies against Muscle Antigens in Facioscapulohumeral Muscular Dystrophy Finds No Disease-Specific Autoantibodies. J Neuromuscul Dis. 2021 May 15;8(5):801–14. doi:10.3233/JND-210653/ASSET/IMAGES/10.3233_JND-210653-FIG7.JPG PubMed PMID: 34024774.

25. Daman K, Yan J, Burzenski LM, Kady J, Shultz LD, Brehm A, et al. A human immune/muscle xenograft model of FSHD muscle pathology. bioRxiv. 2023 Nov 17;2023.11.17.567590. doi:10.1101/2023.11.17.567590 PubMed PMID: 38014123.

26. Banerji CRS, Greco A, Joosten LAB, Van Engelen BGM, Zammit PS. The FSHD muscle-blood biomarker: a circulating transcriptomic biomarker for clinical severity in facioscapulohumeral muscular dystrophy. Brain Commun. 2023;5(5). doi:10.1093/BRAINCOMMS/FCAD221 PubMed PMID: 37731904.

27. Tasca G, Pescatori M, Monforte M, Mirabella M, Iannaccone E, Frusciante R, et al. Different molecular signatures in magnetic resonance imaging-staged facioscapulohumeral muscular dystrophy muscles. PLoS One. 2012 Jun 13;7(6). doi:10.1371/JOURNAL.PONE.0038779 PubMed PMID: 22719944.

28. Wang LH, Friedman SD, Shaw D, Snider L, Wong CJ, Budech CB, et al. MRI-informed muscle biopsies correlate MRI with pathology and DUX4 target gene expression in FSHD. Hum Mol Genet. 2019 Feb 1;28(3):476–86. doi:10.1093/HMG/DDY364 PubMed PMID: 30312408.

29. Wong CJ, Wang LH, Friedman SD, Shaw D, Campbell AE, Budech CB, et al. Longitudinal measures of RNA expression and disease activity in FSHD muscle biopsies. Hum Mol Genet. 2020 Apr 15;29(6):1030–44. doi:10.1093/HMG/DDAA031 PubMed PMID: 32083293.

30. Jones TI, Chew GL, Barraza-Flores P, Schreier S, Ramirez M, Wuebbles RD, et al. Transgenic mice expressing tunable levels of DUX4 develop characteristic facioscapulohumeral muscular dystrophy-like pathophysiology ranging in severity. Skelet Muscle. 2020 Apr 11;10(1). doi:10.1186/S13395-020-00227-4 PubMed PMID: 32278354.

31. van den Heuvel A, Lassche S, Mul K, Greco A, San León Granado D, Heerschap A, et al. Facioscapulohumeral dystrophy transcriptome signatures correlate with different stages of disease and are marked by different MRI biomarkers. Sci Rep. 2022 Dec 1;12(1). doi:10.1038/S41598-022-04817-8 PubMed PMID: 35082321.

32. Sohn S, Reid S, Bowen M, Hudon C, Corbex E, Morel B, et al. AAV-shDUX4 provides short-term benefits but limited long-term efficacy in a DUX4 mouse model of FSHD. Mol Ther Methods Clin Dev. 2025 Sep 11;33(3). doi:10.1016/J.OMTM.2025.101534 PubMed PMID: 40799204.

33. Sohn S, Reid S, Bowen M, Corbex E, Le Gall L, Sidlauskaite E, et al. Molecular, Histological, and Functional Changes in Acta1-MCM;FLExDUX4/+ Mice. Int J Mol Sci. 2024 Nov 1;25(21). doi:10.3390/IJMS252111377 PubMed PMID: 39518930.

34. Lu-Nguyen N, Snowden S, Popplewell L, Malerba A. Systemic Pharmacotherapeutic Treatment of the ACTA1-MCM/FLExDUX4 Preclinical Mouse Model of FSHD. Int J Mol Sci. 2024 Jul 1;25(13). doi:10.3390/IJMS25136994 PubMed PMID: 39000102.

35. Nunes AM, Ramirez MM, Garcia-Collazo E, Jones TI, Jones PL. Muscle eosinophilia is a hallmark of chronic disease in facioscapulohumeral muscular dystrophy. Hum Mol Genet. 2024 May 15;33(10):872–83. doi:10.1093/HMG/DDAE019 PubMed PMID: 38340007.

36. Kakimoto T, Ogasawara A, Ishikawa K, Kurita T, Yoshida K, Harada S, et al. A Systemically Administered Unconjugated Antisense Oligonucleotide Targeting DUX4 Improves Muscular Injury and Motor Function in FSHD Model Mice. Biomedicines. 2023 Sep 1;11(9). doi:10.3390/BIOMEDICINES11092339 PubMed PMID: 37760780.

37. Lu-Nguyen N, Malerba A, Antoni Pineda M, Dickson G, Popplewell L. Improving Molecular and Histopathology in Diaphragm Muscle of the Double Transgenic ACTA1-MCM/FLExDUX4 Mouse Model of FSHD with Systemic Antisense Therapy. Hum Gene Ther. 2022 Sep 1;33(17–18):923–35. doi:10.1089/HUM.2021.251 PubMed PMID: 35078334.

38. Lu-Nguyen N, Dickson G, Malerba A, Popplewell L. Long-Term Systemic Treatment of a Mouse Model Displaying Chronic FSHD-like Pathology with Antisense Therapeutics That Inhibit DUX4 Expression. Biomedicines. 2022 Jul 1;10(7). doi:10.3390/BIOMEDICINES10071623 PubMed PMID: 35884928.

39. Bouwman LF, den Hamer B, van den Heuvel A, Franken M, Jackson M, Dwyer CA, et al. Systemic delivery of a DUX4-targeting antisense oligonucleotide to treat facioscapulohumeral muscular dystrophy. Mol Ther Nucleic Acids. 2021 Dec 3;26:813–27. doi:10.1016/J.OMTN.2021.09.010 PubMed PMID: 34729250.

40. Lu-Nguyen N, Malerba A, Herath S, Dickson G, Popplewell L. Systemic antisense therapeutics inhibiting DUX4 expression ameliorates FSHD-like pathology in an FSHD mouse model. Hum Mol Genet. 2021 Aug 1;30(15):1398–412. doi:10.1093/HMG/DDAB136 PubMed PMID: 33987655.

41. Lim KRQ, Bittel A, Maruyama R, Echigoya Y, Nguyen Q, Huang Y, et al. DUX4 Transcript Knockdown with Antisense 2’-O-Methoxyethyl Gapmers for the Treatment of Facioscapulohumeral Muscular Dystrophy. Mol Ther. 2021 Feb 3;29(2):848–58. doi:10.1016/J.YMTHE.2020.10.010 PubMed PMID: 33068777.

42. Jones T, Jones PL. A cre-inducible DUX4 transgenic mouse model for investigating facioscapulohumeral muscular dystrophy. PLoS One. 2018 Feb 1;13(2). doi:10.1371/JOURNAL.PONE.0192657 PubMed PMID: 29415061.

43. Gros M, Nunes AM, Daoudlarian D, Pini J, Martinuzzi E, Barbosa S, et al. Identification of Serum Interleukin 6 Levels as a Disease Severity Biomarker in Facioscapulohumeral Muscular Dystrophy. J Neuromuscul Dis. 2022 Aug 24;9(1):83–93. doi:10.3233/JND-210711 PubMed PMID: 34459413.

44. Shadle SC, Bennett SR, Wong CJ, Karreman NA, Campbell AE, van der Maarel SM, et al. DUX4-induced bidirectional HSATII satellite repeat transcripts form intranuclear double-stranded RNA foci in human cell models of FSHD. Hum Mol Genet. 2019;28(23). doi:10.1093/hmg/ddz242

45. Shadle SC, Zhong JW, Campbell AE, Conerly ML, Jagannathan S, Wong CJ, et al. DUX4-induced dsRNA and MYC mRNA stabilization activate apoptotic pathways in human cell models of facioscapulohumeral dystrophy. PLoS Genet. 2017;13(3). doi:10.1371/journal.pgen.1006658

46. Rosenberg L, Vabret N. Viral mimicry in cancer therapy. Trends Cancer. 2025 Dec 1;11(12). doi:10.1016/J.TRECAN.2025.08.010 PubMed PMID: 40987678.

47. Chen R, Ishak CA, Carvalho DD de. Endogenous retroelements and the viral mimicry response in cancer therapy and cellular homeostasis. Cancer Discovery. 2021. doi:10.1158/2159-8290.CD-21-0506

48. Contreras O, Rossi FMV, Theret M. Origins, potency, and heterogeneity of skeletal muscle fibro-adipogenic progenitors-time for new definitions. Skelet Muscle. 2021 Dec 1;11(1). doi:10.1186/S13395-021-00265-6 PubMed PMID: 34210364.

49. Wei X, Nicoletti C, Puri PL. Fibro-Adipogenic Progenitors: Versatile keepers of skeletal muscle homeostasis, beyond the response to myotrauma. Semin Cell Dev Biol. 2021 Nov 1;119:23–31. doi:10.1016/J.SEMCDB.2021.07.013 PubMed PMID: 34332886.

50. Singh P, Chazaud B. Benefits and pathologies associated with the inflammatory response. Exp Cell Res. 2021 Dec 1;409(1). doi:10.1016/J.YEXCR.2021.112905 PubMed PMID: 34736921.

51. Coulis G, Jaime D, Guerrero-Juarez C, Kastenschmidt JM, Farahat PK, Nguyen Q, et al. Single-cell and spatial transcriptomics identify a macrophage population associated with skeletal muscle fibrosis. Sci Adv. 2023 Jul 7;9(27). doi:10.1126/SCIADV.ADD9984/SUPPL_FILE/SCIADV.ADD9984_TABLE_S1.ZIP PubMed PMID: 37418531.

52. Pinal-Fernandez I, Casal-Dominguez M, Derfoul A, Pak K, Plotz P, Miller FW, et al. Identification of distinctive interferon gene signatures in different types of myositis. Neurology. 2019 Sep 17;93(12):e1193. doi:10.1212/WNL.0000000000008128 PubMed PMID: 31434690.

53. Pini J, Martinuzzi E, Dhifallah S, Slioui A, Puma A, Villa L, et al. Interleukin-6 as a Key Biomarker in Facioscapulohumeral Dystrophy: Evidence From Longitudinal Analyses. Ann Clin Transl Neurol. 2025. doi:10.1002/ACN3.70210 PubMed PMID: 41058127.

54. Greco A, Mul K, Jaeger MH, Dos Santos JC, Koenen H, De Jong L, et al. IL-6 and TNF are Potential Inflammatory Biomarkers in Facioscapulohumeral Muscular Dystrophy. J Neuromuscul Dis. 2024 Mar 5;11(2):327–47. doi:10.3233/JND-230063 PubMed PMID: 38250782.

55. Jung U, Wei E, Ahsan H, Mitanoska A, Chen K, Kyba M, et al. Matrix metalloproteinases are hallmark early biomarkers and therapeutic targets in FSHD. JCI Insight. 2025 Nov 10;10(21). doi:10.1172/JCI.INSIGHT.195104 PubMed PMID: 40966415.

56. Engquist EN, Greco A, Joosten LAB, van Engelen BGM, Zammit PS, Banerji CRS. FSHD muscle shows perturbation in fibroadipogenic progenitor cells, mitochondrial function and alternative splicing independently of inflammation. Hum Mol Genet. 2024 Jan 15;33(2):182–97. doi:10.1093/HMG/DDAD175 PubMed PMID: 37856562.

57. Di Pietro L, Giacalone F, Ragozzino E, Saccone V, Tiberio F, De Bardi M, et al. Non-myogenic mesenchymal cells contribute to muscle degeneration in facioscapulohumeral muscular dystrophy patients. Cell Death Dis. 2022 Sep 1;13(9). doi:10.1038/S41419-022-05233-6 PubMed PMID: 36114172.

58. Bosnakovski D, Oyler D, Mitanoska A, Douglas M, Ener ET, Shams AS, et al. Persistent Fibroadipogenic Progenitor Expansion Following Transient DUX4 Expression Provokes a Profibrotic State in a Mouse Model for FSHD. Int J Mol Sci. 2022 Feb 1;23(4). doi:10.3390/IJMS23041983 PubMed PMID: 35216102.

59. Bosnakovski D, Shams AS, Yuan C, da Silva MT, Ener ET, Baumann CW, et al. Transcriptional and cytopathological hallmarks of FSHD in chronic DUX4-expressing mice. J Clin Invest. 2020 May 1;130(5):2465–77. doi:10.1172/JCI133303 PubMed PMID: 32250341.

60. Bosnakovski D, Chan SSK, Recht OO, Hartweck LM, Gustafson CJ, Athman LL, et al. Muscle pathology from stochastic low level DUX4 expression in an FSHD mouse model. Nat Commun. 2017 Dec 1;8(1). doi:10.1038/S41467-017-00730-1 PubMed PMID: 28916757.

61. Lemmers RJLF, Van Der Vliet PJ, Klooster R, Sacconi S, Camaño P, Dauwerse JG, et al. A unifying genetic model for facioscapulohumeral muscular dystrophy. Science. 2010 Sep 24;329(5999):1650–3. doi:10.1126/SCIENCE.1189044 PubMed PMID: 20724583.

62. Tihaya MS, Mul K, Balog J, de Greef JC, Tapscott SJ, Tawil R, et al. Facioscapulohumeral muscular dystrophy: the road to targeted therapies. Nat Rev Neurol. 2023 Feb 1;19(2):91–108. doi:10.1038/S41582-022-00762-2 PubMed PMID: 36627512.

63. Ferguson MR, Poliachik SL, Budech CB, Gove NE, Carter GT, Wang LH, et al. MRI change metrics of facioscapulohumeral muscular dystrophy: Stir and T1. Muscle Nerve. 2018 Jun 1;57(6):905–12. doi:10.1002/MUS.26038 PubMed PMID: 29236297.

64. Tasca G, Monforte M, Ottaviani P, Pelliccioni M, Frusciante R, Laschena F, et al. Magnetic resonance imaging in a large cohort of facioscapulohumeral muscular dystrophy patients: Pattern refinement and implications for clinical trials. Ann Neurol. 2016 May 1;79(5):854–64. doi:10.1002/ANA.24640

65. Monforte M, Laschena F, Ottaviani P, Bagnato MR, Pichiecchio A, Tasca G, et al. Tracking muscle wasting and disease activity in facioscapulohumeral muscular dystrophy by qualitative longitudinal imaging. J Cachexia Sarcopenia Muscle. 2019 Dec 1;10(6):1258–65. doi:10.1002/JCSM.12473 PubMed PMID: 31668022.

66. Friedman SD, Poliachik SL, Otto RK, Carter GT, Budech CB, Bird TD, et al. Longitudinal features of stir bright signal in FSHD1. Muscle Nerve. 2014 Feb;49(2):257–60. doi:10.1002/MUS.23911 PubMed PMID: 23720194.

67. Paoletti M, Monforte M, Barzaghi L, Tasca G, Bergsland N, Faggioli A, et al. Natural history of facioscapulohumeral muscular dystrophy evaluated by multiparametric quantitative MRI: a prospective cohort study. J Neurol. 2025 Apr 1;272(4):306-. doi:10.1007/S00415-025-13062-8/TABLES/4 PubMed PMID: 40172709.

68. Bosnakovski D, Xu Z, Ji Gang E, Galindo CL, Liu M, Simsek T, et al. An isogenetic myoblast expression screen identifies DUX4-mediated FSHD-associated molecular pathologies. EMBO Journal. 2008;27(20). doi:10.1038/emboj.2008.201

69. Knopp P, Krom YD, Banerji CRS, Panamarova M, Moyle LA, den Hamer B, et al. DUX4 induces a transcriptome more characteristic of a less-differentiated cell state and inhibits myogenesis. J Cell Sci. 2016;129(20):3816–31. doi:10.1242/JCS.180372 PubMed PMID: 27744317.

70. Bosnakovski D, Toso EA, Hartweck LM, Magli A, Lee HA, Thompson ER, et al. The DUX4 homeodomains mediate inhibition of myogenesis and are functionally exchangeable with the Pax7 homeodomain. J Cell Sci. 2017;130(21):3685–97. doi:10.1242/JCS.205427 PubMed PMID: 28935672.

71. Abad C, Pinal-Fernandez I, Guillou C, Bourdenet G, Drouot L, Cosette P, et al. IFNγ causes mitochondrial dysfunction and oxidative stress in myositis. Nat Commun. 2024 Dec 1;15(1). doi:10.1038/S41467-024-49460-1, PubMed PMID: 38926363.

72. Casal-Dominguez M, Pinal-Fernandez I, Pak K, Muñoz-Braceras S, Milisenda JC, Torres-Ruiz J, et al. Coordinated local RNA overexpression of complement induced by interferon gamma in myositis. Sci Rep. 2023 Dec 1;13(1). doi:10.1038/S41598-023-28838-Z, PubMed PMID: 36739295.

73. Ripoche J, Mitchell JA, Erdei A, Madin C, Moffatt B, Mokoena T, et al. Interferon γ induces synthesis of complement alternative pathway proteins by human endothelial cells in culture. Journal of Experimental Medicine. 1988 Nov 1;168(5):1917–22. doi:10.1084/JEM.168.5.1917, PubMed PMID: 2972796.

74. Gerritsma JSJ, Gerritsen AF, De Ley M, Van Es LA, Daha MR. Interferon-γ induces biosynthesis of complement components C2, C4 and factor H by human proximal tubular epithelial cells. Cytokine. 1997;9(4):276–83. doi:10.1006/cyto.1996.0164 PubMed PMID: 9112336.

75. Mitchell TJ, Naughton M, Norsworthy P, Davies KA, Walport MJ, Morley BJ. IFN-gamma up-regulates expression of the complement components C3 and C4 by stabilization of mRNA. J Immunol. 1996 Jun 1;156(11):4429–34. PubMed PMID: 8666817.

76. Barnum SR, Jones JL, Benveniste EN. Interferon-gamma regulation of C3 gene expression in human astroglioma cells. J Neuroimmunol. 1992;38(3):275–82. doi:10.1016/0165-5728(92)90020-L, PubMed PMID: 1601980.

77. Adhikari A, Cobb B, Eddington S, Becerra N, Kohli P, Pond A, et al. IFN-γ and CIITA modulate IL-6 expression in skeletal muscle. Cytokine X. 2020;2(2). doi:10.1016/j.cytox.2020.100023

78. Sanceau J, Wijdenes J, Revel M, Wietzerbin J. IL-6 and IL-6 receptor modulation by IFN-gamma and tumor necrosis factor-alpha in human monocytic cell line (THP-1). Priming effect of IFN-gamma. J Immunol. 1991 Oct 15;147(8):2630–7. PubMed PMID: 1918983.

79. Yoshida S, Yamada S, Yokose K, Matsumoto H, Fujita Y, Asano T, et al. Interferon-γ induces interleukin-6 production by neutrophils via the Janus kinase (JAK)-signal transducer and activator of transcription (STAT) pathway. BMC Res Notes. 2021 Dec 1;14(1). doi:10.1186/S13104-021-05860-W, PubMed PMID: 34895310.

80. Zheng D, Wondergem A, Kloet S, Willemsen I, Balog J, Tapscott SJ, et al. snRNA-seq analysis in multinucleated myogenic FSHD cells identifies heterogeneous FSHD transcriptome signatures associated with embryonic-like program activation and oxidative stress-induced apoptosis. Hum Mol Genet. 2024 Feb 1;33(3):284–98. doi:10.1093/HMG/DDAD186 PubMed PMID: 37934801.

81. Karpukhina A, Galkin I, Ma Y, Dib C, Zinovkin R, Pletjushkina O, et al. Analysis of genes regulated by DUX4 via oxidative stress reveals potential therapeutic targets for treatment of facioscapulohumeral dystrophy. Redox Biol. 2021 Jul;43:102008. doi:10.1016/j.redox.2021.102008 PubMed PMID: 34030118.

82. Dmitriev P, Saada YB, Dib C, Ansseau E, Barat A, Hamade A, et al. DUX4-induced constitutive DNA damage and oxidative stress contribute to aberrant differentiation of myoblasts from FSHD patients. Free Radic Biol Med. 2016;99. doi:10.1016/j.freeradbiomed.2016.08.007

83. Passerieux E, Hayot M, Jaussent A, Carnac G, Gouzi F, Pillard F, et al. Effects of vitamin C, vitamin E, zinc gluconate, and selenomethionine supplementation on muscle function and oxidative stress biomarkers in patients with facioscapulohumeral dystrophy: a double-blind randomized controlled clinical trial. Free Radic Biol Med. 2015 Apr;81:158–69. doi:10.1016/j.freeradbiomed.2014.09.014 PubMed PMID: 25246239.

84. Turki A, Hayot M, Carnac G, Pillard F, Passerieux E, Bommart S, et al. Functional muscle impairment in facioscapulohumeral muscular dystrophy is correlated with oxidative stress and mitochondrial dysfunction. Free Radic Biol Med. 2012 Sep 1;53(5):1068–79. doi:10.1016/j.freeradbiomed.2012.06.041 PubMed PMID: 22796148.

85. Barro M, Carnac G, Flavier S, Mercier J, Vassetzky Y, Laoudj-Chenivesse D. Myoblasts from affected and non-affected FSHD muscles exhibit morphological differentiation defects. J Cell Mol Med. 2010 Jan;14(1–2):275–89. doi:10.1111/j.1582-4934.2008.00368.x PubMed PMID: 18505476.

86. Winokur ST, Barrett K, Martin JH, Forrester JR, Simon M, Tawil R, et al. Facioscapulohumeral muscular dystrophy (FSHD) myoblasts demonstrate increased susceptibility to oxidative stress. Neuromuscul Disord. 2003 May;13(4):322–33. PubMed PMID: 12868502.

87. Laoudj-Chenivesse D, Carnac G, Bisbal C, Hugon G, Bouillot S, Desnuelle C, et al. Increased levels of adenine nucleotide translocator 1 protein and response to oxidative stress are early events in facioscapulohumeral muscular dystrophy muscle. J Mol Med. 2005 Mar;83(3):216–24. doi:10.1007/S00109-004-0583-7, PubMed PMID: 15551024.

88. Macaione V, Aguennouz M, Rodolico C, Mazzeo A, Patti A, Cannistraci E, et al. RAGE-NF-κB pathway activation in response to oxidative stress in facioscapulohumeral muscular dystrophy. Acta Neurol Scand. 2007 Feb;115(2):115–21. doi:10.1111/J.1600-0404.2006.00724.X, PubMed PMID: 17212615.

89. Sasaki-Honda M, Jonouchi T, Arai M, Hotta A, Mitsuhashi S, Nishino I, et al. A patient-derived iPSC model revealed oxidative stress increases facioscapulohumeral muscular dystrophy-causative DUX4. Hum Mol Genet. 2018 Dec 1;27(23):4024–35. doi:10.1093/HMG/DDY293, PubMed PMID: 30107443.

90. Heher P, Ganassi M, Weidinger A, Engquist EN, Pruller J, Nguyen TH, et al. Interplay between mitochondrial reactive oxygen species, oxidative stress and hypoxic adaptation in facioscapulohumeral muscular dystrophy: Metabolic stress as potential therapeutic target. Redox Biol. 2022 May 1;51. doi:10.1016/j.redox.2022.102251 PubMed PMID: 35248827.

91. Wilson VD, Bommart S, Passerieux E, Thomas C, Pincemail J, Picot MC, et al. Muscle strength, quantity and quality and muscle fat quantity and their association with oxidative stress in patients with facioscapulohumeral muscular dystrophy: Effect of antioxidant supplementation. Free Radic Biol Med. 2024 Jul 1;219:112–26. doi:10.1016/j.freeradbiomed.2024.04.001 PubMed PMID: 38574978.

92. Moriggi M, Ruggiero L, Torretta E, Zoppi D, Arosio B, Ferri E, et al. Muscle Proteome Analysis of Facioscapulohumeral Dystrophy Patients Reveals a Metabolic Rewiring Promoting Oxidative/Reductive Stress Contributing to the Loss of Muscle Function. Antioxidants. 2024 Nov 1;13(11). doi:10.3390/ANTIOX13111406.

93. Wong CJ, Wang L, Holers VM, Frazer-Abel A, Van Der Maarel SM, Tawil R, et al. Elevated plasma complement components in facioscapulohumeral dystrophy. Hum Mol Genet. 2022 Jun 1;31(11):1821–9. doi:10.1093/HMG/DDAB364, PubMed PMID: 34919696.

94. Rahimov F, King OD, Leung DG, Bibat GM, Emerson CP, Kunkel LM, et al. Transcriptional profiling in facioscapulohumeral muscular dystrophy to identify candidate biomarkers. Proc Natl Acad Sci U S A. 2012 Oct 2;109(40):16234–9. doi:10.1073/PNAS.1209508109, PubMed PMID: 22988124.

95. Spuler S, Engel AG. Unexpected sarcolemmal complement membrane attack complex deposits on nonnecrotic muscle fibers in muscular dystrophies. Neurology. 1998;50(1):41–6. doi:10.1212/WNL.50.1.41, PubMed PMID: 9443455.

96. Wei TT, Zhang MY, Zheng XH, Xie TH, Wang W, Zou J, et al. Interferon-γ induces retinal pigment epithelial cell Ferroptosis by a JAK1-2/STAT1/SLC7A11 signaling pathway in Age-related Macular Degeneration. FEBS J. 2022 Apr;289(7):1968–83. doi:10.1111/febs.16272 PubMed PMID: 34741776.

97. Cao T, Zhou J, Liu Q, Mao T, Chen B, Wu Q, et al. Interferon-γ induces salivary gland epithelial cell ferroptosis in Sjogren’s syndrome via JAK/STAT1-mediated inhibition of system Xc. Free Radic Biol Med. 2023 Aug 20;205:116–28. doi:10.1016/j.freeradbiomed.2023.05.027 PubMed PMID: 37286044.

98. Huang J, Yu S, Luo J, Luo X, Yang J, Wang X. IFN-γ could induce ferroptosis in keloid fibroblasts by inhibiting the expression of serpine2. Cell Death Discov. 2025 May 5;11(1):217. doi:10.1038/s41420-025-02401-3 PubMed PMID: 40324981.

99. Yu X, Zhu D, Luo B, Kou W, Cheng Y, Zhu Y. IFNγ enhances ferroptosis by increasing JAK-STAT pathway activation to suppress SLCA711 expression in adrenocortical carcinoma. Oncol Rep. 2022 May;47(5). doi:10.3892/or.2022.8308 PubMed PMID: 35322867.

100. Wang X, Yang Y, Zhou X, Yu S, Luo X, Lu L, et al. IFNγ regulates ferroptosis in KFs by inhibiting the expression of SPOCD1 through DNMT3A. Cell Death Discov. 2025 Jan 16;11(1):9. doi:10.1038/s41420-024-02257-z PubMed PMID: 39820341.

101. Nakamura K, Ortuste-Quiroga HP, Horii N, Fujimaki S, Moroishi T, Nakayama KI, et al. Iron supplementation alleviates pathologies in a mouse model of facioscapulohumeral muscular dystrophy. J Clin Invest. 2025 Jul 1. doi:10.1172/JCI181881 PubMed PMID: 40591405.

102. Hou C, Periou B, Gervais M, Martin L, Berthier J, Baba-Amer Y, et al. Interferon-γ causes myogenic cell dysfunction and senescence in immune myopathies. Brain. 2025 Aug 1;148(8):2883–98. doi:10.1093/BRAIN/AWAF153 PubMed PMID: 40296760.

103. Corasolla Carregari V, Monforte M, Di Maio G, Pieroni L, Urbani A, Ricci E, et al. Proteomics of Muscle Microdialysates Identifies Potential Circulating Biomarkers in Facioscapulohumeral Muscular Dystrophy. Int J Mol Sci. 2020 Dec 30;22(1). doi:10.3390/IJMS22010290 PubMed PMID: 33396627.

104. Geng LN, Yao Z, Snider L, Fong AP, Cech JN, Young JM, et al. DUX4 Activates Germline Genes, Retroelements, and Immune Mediators: Implications for Facioscapulohumeral Dystrophy. Dev Cell. 2012;22(1). doi:10.1016/j.devcel.2011.11.013

105. Whiddon JL, Langford AT, Wong CJ, Zhong JW, Tapscott SJ. Conservation and innovation in the DUX4-family gene network. Nat Genet. 2017 Jun 1;49(6):935–40. doi:10.1038/NG.3846 PubMed PMID: 28459454.

106. Hendrickson PG, Doráis JA, Grow EJ, Whiddon JL, Lim JW, Wike CL, et al. Conserved roles of mouse DUX and human DUX4 in activating cleavage-stage genes and MERVL/HERVL retrotransposons. Nat Genet. 2017 Jun 1;49(6):925–34. doi:10.1038/NG.3844 PubMed PMID: 28459457.

107. Campolungo D, Salomé M, Biferali B, Tascini AS, Gabellini D. DUX4-r exerts a neomorphic activity that depends on GTF2I in acute lymphoblastic leukemia. Sci Adv. 2023;9(37). doi:10.1126/SCIADV.ADI3771 PubMed PMID: 37713484.

108. Zheng D, van den Heuvel A, Balog J, Willemsen IM, Kloet S, Tapscott SJ, et al. DUX4 activates common and context-specific intergenic transcripts and isoforms. Sci Adv. 2025 May 9;11(19). doi:10.1126/SCIADV.ADT5356 PubMed PMID: 40333962.

109. Sato M, Suemori H, Hata N, Asagiri M, Ogasawara K, Nakao K, et al. Distinct and essential roles of transcription factors IRF-3 and IRF-7 in response to viruses for IFN-alpha/beta gene induction. Immunity. 2000;13(4):539–48. doi:10.1016/S1074-7613(00)00053-4 PubMed PMID: 11070172.

110. Duan T, Du Y, Xing C, Wang HY, Wang RF. Toll-Like Receptor Signaling and Its Role in Cell-Mediated Immunity. Front Immunol. 2022 Mar 3;13. doi:10.3389/FIMMU.2022.812774 PubMed PMID: 35309296.

111. Jagannathan S, Shadle SC, Resnick R, Snider L, Tawil RN, van der Maarel SM, et al. Model systems of DUX4 expression recapitulate the transcriptional profile of FSHD cells. Hum Mol Genet. 2016;25(20):4419–31. doi:10.1093/HMG/DDW271 PubMed PMID: 28171552.

112. Zhao Y, Liang L, Fan Y, Sun S, An L, Shi Z, et al. PPM1B negatively regulates antiviral response via dephosphorylating TBK1. Cell Signal. 2012 Nov;24(11):2197–204. doi:10.1016/J.CELLSIG.2012.06.017 PubMed PMID: 22750291.

113. Gao M, Cheng X, Zhang C, Ma J, Tian X, Wang S, et al. Human-specific activation of the DUX4-SLC34A2 axis by herpesviruses suppresses antiviral innate immunity. mBio. 2025 Dec 10;16(12):e0255425. doi:10.1128/MBIO.02554-25 PubMed PMID: 41211819.

114. Miyashita M, Oshiumi H, Matsumoto M, Seya T. DDX60, a DEXD/H box helicase, is a novel antiviral factor promoting RIG-I-like receptor-mediated signaling. Mol Cell Biol. 2011 Sep 1;31(18):3802–19. doi:10.1128/MCB.01368-10 PubMed PMID: 21791617.

115. Shen Y, Li NL, Wang J, Liu B, Lester S, Li K. TRIM56 is an essential component of the TLR3 antiviral signaling pathway. J Biol Chem. 2012 Oct 19;287(43):36404–13. doi:10.1074/JBC.M112.397075 PubMed PMID: 22948160.

116. Carlin AF, Plummer EM, Vizcarra EA, Sheets N, Joo Y, Tang W, et al. An IRF-3-, IRF-5-, and IRF-7-Independent Pathway of Dengue Viral Resistance Utilizes IRF-1 to Stimulate Type I and II Interferon Responses. Cell Rep. 2017 Nov 7;21(6):1600–12. doi:10.1016/J.CELREP.2017.10.054 PubMed PMID: 29117564.

117. Walker WE, Garcia LF, Palermo PM, Hakim N, Goswami DG, Dwivedi AK, et al. Interferon Regulatory Factor 3 Exacerbates the Severity of COVID-19 in Mice. Crit Care Explor. 2025 Mar 17;7(3):e1225. doi:10.1097/CCE.0000000000001225 PubMed PMID: 40103621.

118. Green DS, Young HA, Valencia JC. Current prospects of type II interferon γ signaling and autoimmunity. J Biol Chem. 2017 Aug 25;292(34):13925–33. doi:10.1074/jbc.R116.774745 PubMed PMID: 28652404.

119. Spens AE, Sutliff NA, Bennett SR, Campbell AE, Tapscott SJ. Human DUX4 and mouse Dux interact with STAT1 and broadly inhibit interferon-stimulated gene induction. Elife. 2023 Apr 24;12. doi:10.7554/ELIFE.82057 PubMed PMID: 37092726.

120. Ramana C V., Gil MP, Han Y, Ransohoff RM, Schreiber RD, Stark GR. Stat1-independent regulation of gene expression in response to IFN-gamma. Proc Natl Acad Sci U S A. 2001 Jun 5;98(12):6674–9. doi:10.1073/PNAS.111164198 PubMed PMID: 11390994.

121. Gil MP, Bohn E, O’Guin AK, Ramana C V., Levine B, Stark GR, et al. Biologic consequences of Stat1-independent IFN signaling. Proc Natl Acad Sci U S A. 2001 Jun 5;98(12):6680–5. doi:10.1073/PNAS.111163898 PubMed PMID: 11390995.

122. Ramana C V., Grammatikakis N, Chernov M, Nguyen H, Goh KC, Williams BRG, et al. Regulation of c-myc expression by IFN-gamma through Stat1-dependent and -independent pathways. EMBO J. 2000 Jan 17;19(2):263–72. doi:10.1093/EMBOJ/19.2.263 PubMed PMID: 10637230.

123. Soler C, Felipe A, García-Manteiga J, Serra M, Guillén-Gómez E, Casado FJ, et al. Interferon-gamma regulates nucleoside transport systems in macrophages through signal transduction and activator of transduction factor 1 (STAT1)-dependent and -independent signalling pathways. Biochem J. 2003 Nov 1;375(Pt 3):777–83. doi:10.1042/BJ20030260 PubMed PMID: 12868960.

124. Matsuzawa T, Kim BH, Shenoy AR, Kamitani S, Miyake M, MacMicking JD. IFN-γ elicits macrophage autophagy via the p38 MAPK signaling pathway. J Immunol. 2012 Jul 15;189(2):813–8. doi:10.4049/JIMMUNOL.1102041 PubMed PMID: 22675202.

125. Pasieka TJ, Cilloniz C, Lu B, Teal TH, Proll SC, Katze MG, et al. Host responses to wild-type and attenuated herpes simplex virus infection in the absence of Stat1. J Virol. 2009 Mar;83(5):2075–87. doi:10.1128/JVI.02007-08 PubMed PMID: 19109391.

126. Ramana C V., Kumar A, Enelow R. Stat1-independent induction of SOCS-3 by interferon-gamma is mediated by sustained activation of Stat3 in mouse embryonic fibroblasts. Biochem Biophys Res Commun. 2005 Feb 18;327(3):727–33. doi:10.1016/J.BBRC.2004.12.074 PubMed PMID: 15649407.

127. Qing Y, Stark GR. Alternative activation of STAT1 and STAT3 in response to interferon-gamma. J Biol Chem. 2004 Oct 1;279(40):41679–85. doi:10.1074/JBC.M406413200 PubMed PMID: 15284232.

128. Walters DM, Antao-Menezes A, Ingram JL, Rice AB, Nyska A, Tani Y, et al. Susceptibility of signal transducer and activator of transcription-1-deficient mice to pulmonary fibrogenesis. Am J Pathol. 2005;167(5):1221–9. doi:10.1016/S0002-9440(10)61210-2 PubMed PMID: 16251407.

129. Soler C, Felipe A, García-Manteiga J, Serra M, Guillén-Gómez E, Casado FJ, et al. Interferon-gamma regulates nucleoside transport systems in macrophages through signal transduction and activator of transduction factor 1 (STAT1)-dependent and -independent signalling pathways. Biochem J. 2003 Nov 1;375(Pt 3):777–83. doi:10.1042/BJ20030260 PubMed PMID: 12868960.

130. O’Donnell LA, Henkins KM, Kulkarni A, Matullo CM, Balachandran S, Pattisapu AK, et al. Interferon gamma induces protective non-canonical signaling pathways in primary neurons. J Neurochem. 2015 Oct 1;135(2):309–22. doi:10.1111/JNC.13250 PubMed PMID: 26190522.

131. Ramana C V., Deberge MP, Kumar A, Alia CS, Durbin JE, Enelow RI. Inflammatory impact of IFN-γ in CD8+ T cell-mediated lung injury is mediated by both Stat1-dependent and - independent pathways. Am J Physiol Lung Cell Mol Physiol. 2015;308(7):L650–7. doi:10.1152/AJPLUNG.00360.2014 PubMed PMID: 25617378.

132. Bittel AJ, Bittel DC, Gordish-Dressman H, Chen YW. Voluntary wheel running improves molecular and functional deficits in a murine model of facioscapulohumeral muscular dystrophy. iScience. 2023 Jan 19;27(1). doi:10.1016/J.ISCI.2023.108632 PubMed PMID: 38188524.

133. Amar D, Lindholm ME, Norrbom J, Wheeler MT, Rivas MA, Ashley EA. Time trajectories in the transcriptomic response to exercise - a meta-analysis. Nature Communications 2021 12:1. 2021 Jun 9;12(1):3471-. doi:10.1038/S41467-021-23579-X PubMed PMID: 34108459.

134. Langston PK, Sun Y, Ryback BA, Mueller AL, Spiegelman BM, Benoist C, et al. Regulatory T cells shield muscle mitochondria from interferon-γ–mediated damage to promote the beneficial effects of exercise. Sci Immunol. 2023 Nov 1;8(89). doi:10.1126/SCIIMMUNOL.ADI5377/SUPPL_FILE/SCIIMMUNOL.ADI5377_MDAR_REPRODUCIBILITY_CHECKLIST.PDF PubMed PMID: 37922340.

135. Zamani A, Salehi I, Alahgholi-Hajibehzad M. Moderate Exercise Enhances the Production of Interferon-γ and Interleukin-12 in Peripheral Blood Mononuclear Cells. Immune Netw. 2017;17(3):186–91. doi:10.4110/IN.2017.17.3.186 PubMed PMID: 28680380.

136. Heidarianpour A, Rezazadeh MV, Zamani A, Heidarianpour A, Rezazadeh MV, Zamani A, et al. Effect of Moderate Exercise on Serum Interferon-Gamma and Interleukin-17 Levels in the Morphine Withdrawal Period. International Journal of High Risk Behaviors and Addiction 2016 5:2. 2016 Jun 1;5(2):e19703. doi:10.5812/IJHRBA.26907

137. Al-Salama ZT. Emapalumab: First Global Approval. Drugs. 2019 Jan 1;79(1):99–103. doi:10.1007/S40265-018-1046-8 PubMed PMID: 30623346.

138. Masarova L, Mascarenhas J, Rampal R, Hu W, Livingston RA, Pemmaraju N. Ten years of experience with ruxolitinib since approval for polycythemia vera: A review of clinical efficacy and safety. Cancer. 2025 Jan 1;131(1). doi:10.1002/CNCR.35661 PubMed PMID: 39616447.

139. Santos M Dos, Gioftsidi S, Backer S, Machado L, Relaix F, Maire P, et al. Extraction and sequencing of single nuclei from murine skeletal muscles. STAR Protoc. 2021 Sep 17;2(3):100694. doi:10.1016/J.XPRO.2021.100694 PubMed PMID: 34382019.

140. Fleming SJ, Chaffin MD, Arduini A, Akkad AD, Banks E, Marioni JC, et al. Unsupervised removal of systematic background noise from droplet-based single-cell experiments using CellBender. Nat Methods. 2023 Sep 1;20(9):1323–35. doi:10.1038/S41592-023-01943-7 PubMed PMID: 37550580.

141. Hao Y, Stuart T, Kowalski MH, Choudhary S, Hoffman P, Hartman A, et al. Dictionary learning for integrative, multimodal and scalable single-cell analysis. Nat Biotechnol. 2024 Feb 1;42(2):293–304. doi:10.1038/S41587-023-01767-Y PubMed PMID: 37231261.

142. Marsh S, Salmon M, Hoffman P, kew24, Pir MS. samuel-marsh/scCustomize: Release 3.2.4 [Internet]. doi:10.5281/ZENODO.17900382

143. Korsunsky I, Millard N, Fan J, Slowikowski K, Zhang F, Wei K, et al. Fast, sensitive and accurate integration of single-cell data with Harmony. Nat Methods. 2019 Dec 1;16(12):1289–96. doi:10.1038/S41592-019-0619-0 PubMed PMID: 31740819.

144. Petrany MJ, Swoboda CO, Sun C, Chetal K, Chen X, Weirauch MT, et al. Single-nucleus RNA-seq identifies transcriptional heterogeneity in multinucleated skeletal myofibers. Nature Communications 2020 11:1. 2020 Dec 11;11(1):6374-. doi:10.1038/s41467-020-20063-w PubMed PMID: 33311464.

145. Ham AS, Lin S, Tse A, Thürkauf M, McGowan TJ, Jörin L, et al. Single-nuclei sequencing of skeletal muscle reveals subsynaptic-specific transcripts involved in neuromuscular junction maintenance. Nature Communications 2025 16:1. 2025 Mar 5;16(1):2220-. doi:10.1038/s41467-025-57487-1 PubMed PMID: 40044687.

146. Dos Santos M, Backer S, Saintpierre B, Izac B, Andrieu M, Letourneur F, et al. Single-nucleus RNA-seq and FISH identify coordinated transcriptional activity in mammalian myofibers. Nature Communications 2020 11:1. 2020 Oct 9;11(1):5102-. doi:10.1038/s41467-020-18789-8 PubMed PMID: 33037211.

147. Oprescu SN, Yue F, Qiu J, Brito LF, Kuang S. Temporal Dynamics and Heterogeneity of Cell Populations during Skeletal Muscle Regeneration. iScience. 2020 Apr 24;23(4). doi:10.1016/J.ISCI.2020.100993 PubMed PMID: 32248062.

148. De Micheli AJ, Laurilliard EJ, Heinke CL, Ravichandran H, Fraczek P, Soueid-Baumgarten S, et al. Single-Cell Analysis of the Muscle Stem Cell Hierarchy Identifies Heterotypic Communication Signals Involved in Skeletal Muscle Regeneration. Cell Rep. 2020 Mar 10;30(10):3583–3595.e5. doi:10.1016/J.CELREP.2020.02.067 PubMed PMID: 32160558.

149. Hackert NS, Radtke FA, Exner T, Lorenz HM, Müller-Tidow C, Nigrovic PA, et al. Human and mouse neutrophils share core transcriptional programs in both homeostatic and inflamed contexts. Nature Communications 2023 14:1. 2023 Dec 8;14(1):8133-. doi:10.1038/s41467-023-43573-9 PubMed PMID: 38065997.

150. Gurtner A, Borrelli C, Gonzalez-Perez I, Bach K, Acar IE, Núñez NG, et al. Active eosinophils regulate host defence and immune responses in colitis. Nature 2022 615:7950. 2022 Dec 12;615(7950):151–7. doi:10.1038/s41586-022-05628-7 PubMed PMID: 36509106.

151. Germain PL, Lun A, Macnair W, Robinson MD. Doublet identification in single-cell sequencing data using scDblFinder. F1000Research 2021 10:979. 2021 Sep 28;10:979. doi:10.12688/f1000research.73600.1 PubMed PMID: 35814628.

152. Howe DG, Blake JA, Bradford YM, Bult CJ, Calvi BR, Engel SR, et al. Model organism data evolving in support of translational medicine. Lab Anim (NY). 2018 Oct 1;47(10):277–89. doi:10.1038/S41684-018-0150-4 PubMed PMID: 30224793.

153. Tirosh I, Izar B, Prakadan SM, Wadsworth MH, Treacy D, Trombetta JJ, et al. Dissecting the multicellular ecosystem of metastatic melanoma by single-cell RNA-seq. Science. 2016 Apr 8;352(6282):189–96. doi:10.1126/SCIENCE.AAD0501 PubMed PMID: 27124452.

154. Wickham H. Data Analysis [Internet]. 2016;189–201. doi:10.1007/978-3-319-24277-4_9

155. Parks B, Greenleaf W. Scalable high-performance single cell data analysis with BPCells. bioRxiv. 2025 Apr 1;2025.03.27.645853. doi:10.1101/2025.03.27.645853

156. Andreatta M, Carmona SJ. UCell: Robust and scalable single-cell gene signature scoring. Comput Struct Biotechnol J. 2021 Jan 1;19:3796–8. doi:10.1016/J.CSBJ.2021.06.043 PubMed PMID: 34285779.

157. Robinson MD, McCarthy DJ, Smyth GK. edgeR: a Bioconductor package for differential expression analysis of digital gene expression data. Bioinformatics. 2010 Nov 11;26(1):139–40. doi:10.1093/BIOINFORMATICS/BTP616 PubMed PMID: 19910308.

158. Liberzon A, Birger C, Thorvaldsdóttir H, Ghandi M, Mesirov JP, Tamayo P. The Molecular Signatures Database (MSigDB) hallmark gene set collection. Cell Syst. 2015 Dec 23;1(6):417–25. doi:10.1016/J.CELS.2015.12.004 PubMed PMID: 26771021.

159. Castanza AS, Recla JM, Eby D, Thorvaldsdóttir H, Bult CJ, Mesirov JP. Extending support for mouse data in the Molecular Signatures Database (MSigDB). Nat Methods. 2023 Nov 1;20(11):1619–20. doi:10.1038/S41592-023-02014-7 PubMed PMID: 37704782.

160. Chen EY, Tan CM, Kou Y, Duan Q, Wang Z, Meirelles G V., et al. Enrichr: interactive and collaborative HTML5 gene list enrichment analysis tool. BMC Bioinformatics. 2013 Apr 15;14. doi:10.1186/1471-2105-14-128 PubMed PMID: 23586463.

161. Kuleshov M V., Jones MR, Rouillard AD, Fernandez NF, Duan Q, Wang Z, et al. Enrichr: a comprehensive gene set enrichment analysis web server 2016 update. Nucleic Acids Res. 2016 Jul 8;44(W1):W90–7. doi:10.1093/NAR/GKW377 PubMed PMID: 27141961.

162. Xie Z, Bailey A, Kuleshov M V., Clarke DJB, Evangelista JE, Jenkins SL, et al. Gene Set Knowledge Discovery with Enrichr. Curr Protoc. 2021 Mar 1;1(3). doi:10.1002/CPZ1.90 PubMed PMID: 33780170.

163. Subramanian A, Tamayo P, Mootha VK, Mukherjee S, Ebert BL, Gillette MA, et al. Gene set enrichment analysis: A knowledge-based approach for interpreting genome-wide expression profiles. Proc Natl Acad Sci U S A. 2005 Oct 25;102(43):15545–50. doi:10.1073/PNAS.0506580102/SUPPL_FILE/06580FIG7.JPG PubMed PMID: 16199517.

164. Biferali B, Bianconi V, Perez DF, Kronawitter SP, Marullo F, Maggio R, et al. Prdm16-mediated H3K9 methylation controls fibro-adipogenic progenitors identity during skeletal muscle repair. Sci Adv. 2021;7(23). doi:10.1126/sciadv.abd9371

165. de Faria IJS, Imler JL, Marques JT. Protocol for the analysis of double-stranded RNAs in virus-infected insect cells using anti-dsRNA antibodies. STAR Protoc. 2023 Mar 17;4(1):102033. doi:10.1016/J.XPRO.2022.102033 PubMed PMID: 36853733.

166. Bankhead P, Loughrey MB, Fernández JA, Dombrowski Y, McArt DG, Dunne PD, et al. QuPath: Open source software for digital pathology image analysis. Scientific Reports 2017 7:1. 2017 Dec 4;7(1):16878-. doi:10.1038/s41598-017-17204-5 PubMed PMID: 29203879.

