## Supplementary Figures for "The interferon-gamma response pathway is a targetable driver of FSHD muscular dystrophy progression"

### Supplementary Materials

**Fig. S1.**

#### Supplementary 1

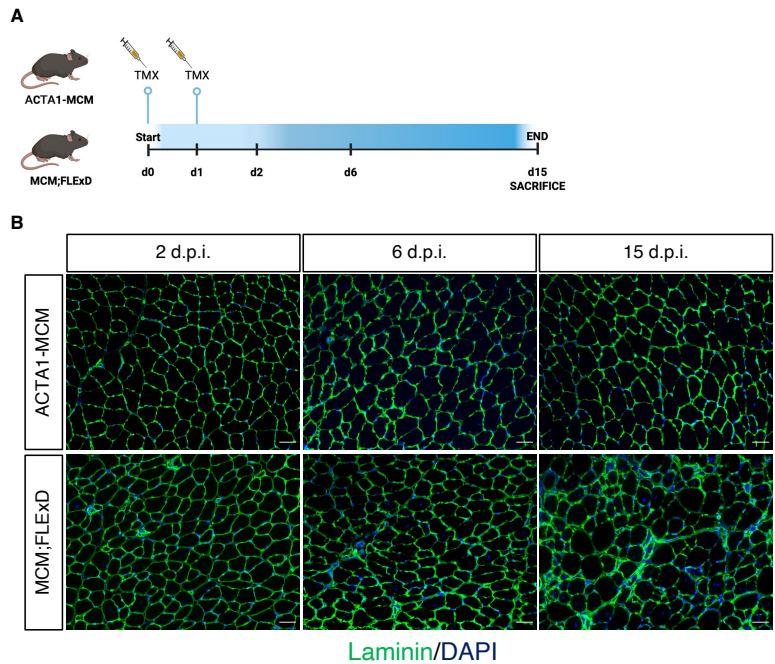

**Figure S1. TMX regimen for ACTA1-MCM and MCM;FLEXD mice**

(A) Schematic representation of the applied TMX regimen for MCM;FLEXD and ACTA1-MCM mice. Mice were treated with 5mg/kg of TMX in two consecutive days (d0 and d1) and then sacrificed upon 2-6-15 d.p.i.. Created with BioRender.com. (B) Representative images of laminin (green) immunofluorescence at 2-6-15 d.p.i. on TA from ACTA1-MCM and MCM;FLEXD mice. Nuclei stained with DAPI (blue). Scale bar: 50μm.

Fig. S2.

#### Supplementary 2

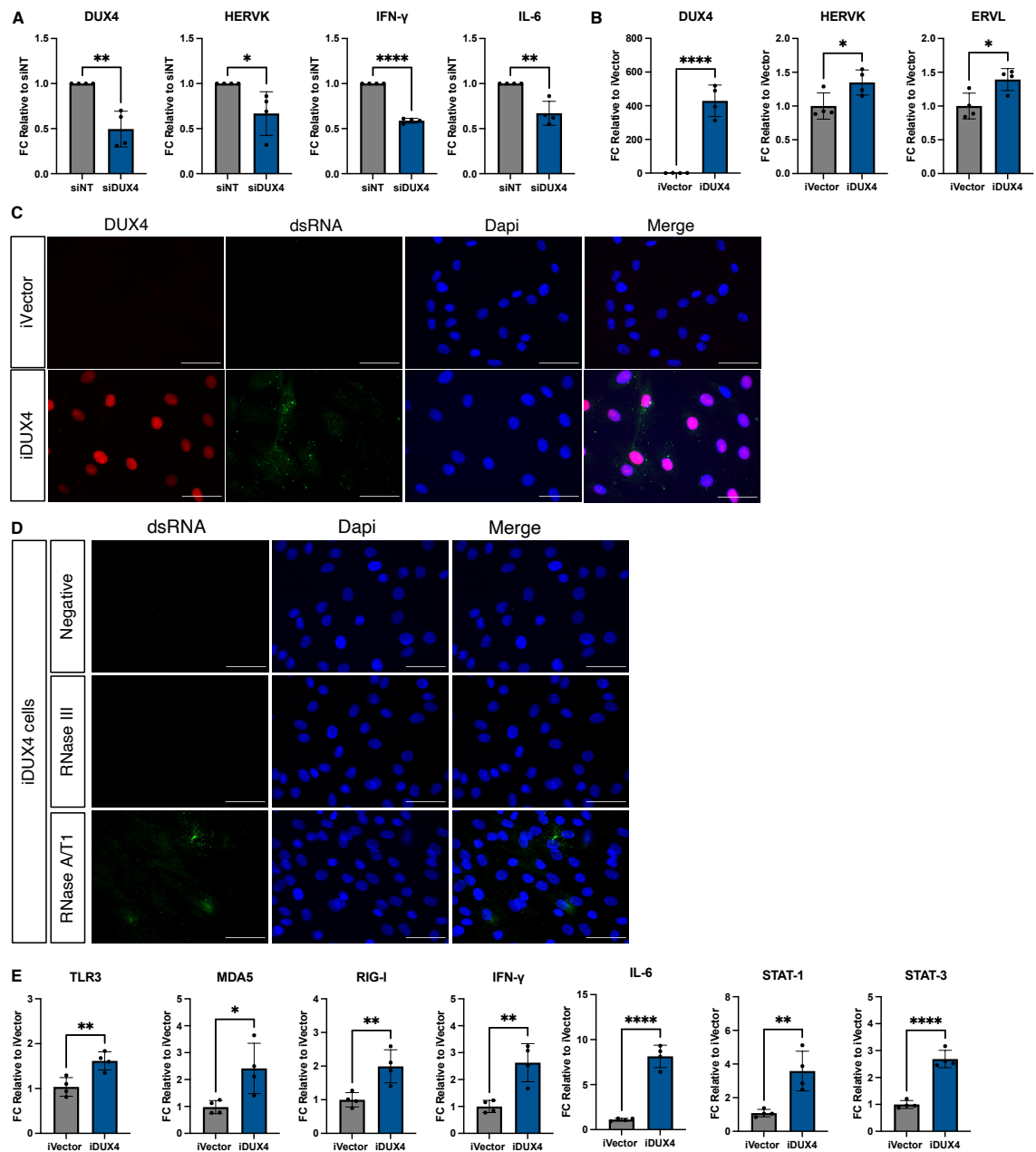

**Figure S2. DUX4 is required and sufficient to activate a viral mimicry-like response**

(A) Expression of *DUX4* and the indicated viral mimicry pathway genes evaluated by RT-qPCR in differentiated human primary FSHD muscle cells treated with control (siNT) or *DUX4* (siDUX4) siRNAs. Values expressed as fold change relative to siNT cells. $n=4$. Statistical significance assessed by Unpaired t-test, \*$p<0.05$; \*\*$p<0.01$; \*\*\*\*$p<0.0001$. (B) Expression of *DUX4* and its indicated repeat target genes evaluated by RT-qPCR in human muscle cells expressing the empty vector (iVector) or *DUX4* (iDUX4) upon doxycycline induction. Values expressed as fold change relative to iVector cells. n=4. Statistical significance assessed by Unpaired t-test, \*p<0.05; \*\*\*\*p<0.0001. (C) Representative images of dsRNA (green) and DUX4 (red) staining in cells in B. Nuclei stained with DAPI (blue). Scale bar: 50µm. (D) Representative images of iDUX4 cells stained just with the secondary antibody (negative) or stained for dsRNA (green) after pre-treatment with RNase III (to digests dsRNA) or RNase A/T1 (to digests single-stranded RNA). Fluorescence is reduced only after RNase III treatment confirming that the signal is specific for dsRNA. Nuclei stained with DAPI (blue). Scale bar: 50µm. (E) Expression the indicated viral mimicry pathway genes evaluated by RT-qPCR in iVector or iDUX4 cells. Values expressed as fold change relative to iVector cells. n=4. Statistical significance assessed by Unpaired t-test, \*p<0.05; \*\*p<0.01; \*\*\*\*p<0.0001.

**Fig. S3.**

**Supplementary 3**

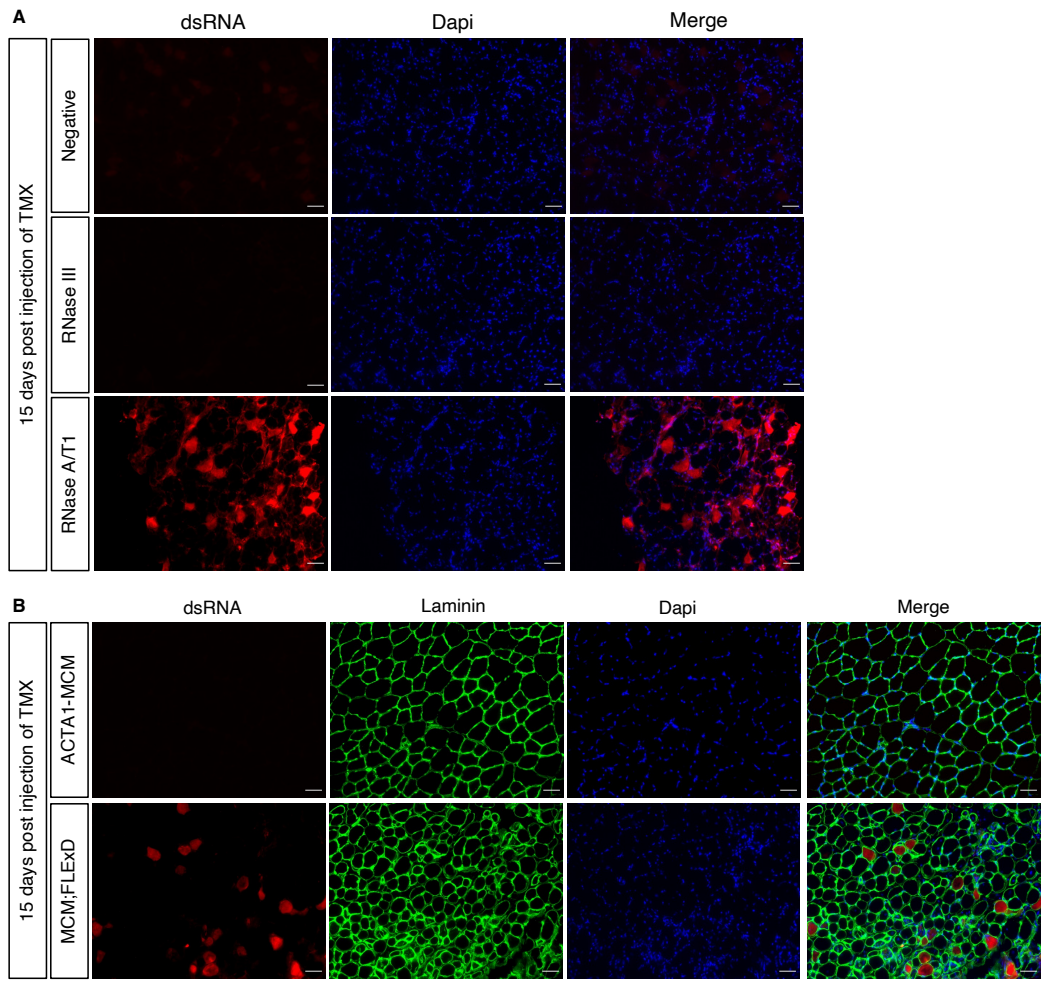

**Figure S3. dsRNA accumulation specifically in MCM;FLEXD skeletal muscle fibers**

(A) Representative images of TA from MCM;FLEXD stained just with the secondary antibody (negative) or stained for dsRNA (red) after pre-treatment with RNase III (to digests dsRNA) or RNase A/T1 (to digests single-stranded RNA). Fluorescence is reduced only after RNase III treatment confirming that the signal is specific for dsRNA. Scale bar: 50μm. (B) Representative images of TA from MCM;FLEXD at 15 d.p.i. stained for dsRNA (red), laminin (green) and DAPI (blue). Scale bar: 50μm.

**Fig. S4.**

**Supplementary 4**

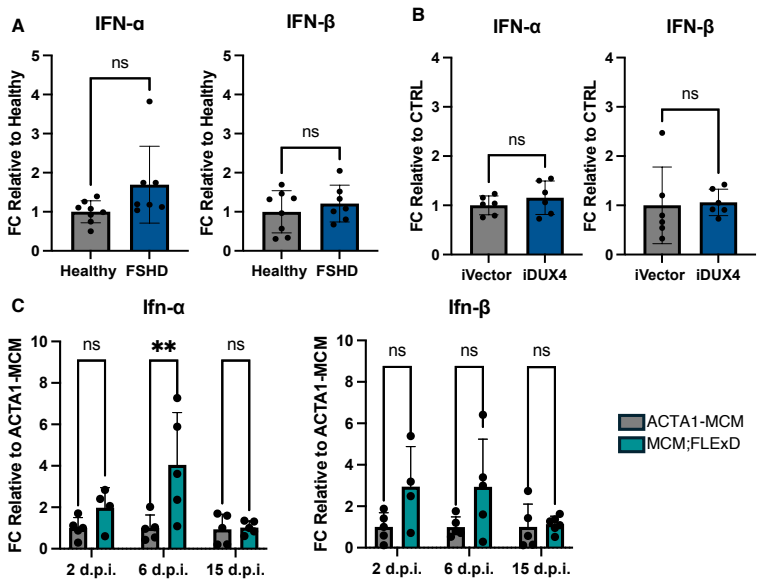

**Figure S4. *IFN-α* and *IFN-β* are not consistently activated in FSHD models**

(A) Expression of *IFN-α* and *IFN-β* genes evaluated by RT-qPCR in differentiated human primary muscle cells from healthy donors (n=8) and FSHD patients (n=7). Values expressed as fold change relative to healthy donor cells. Statistical significance assessed by Unpaired t-test. (B) Expression of *IFN-α* and *IFN-β* genes evaluated by RT-qPCR in iVector and iDUX4 human muscle cells upon doxycycline induction. Values expressed as fold change relative to iVector cells. n=6. Statistical significance assessed by Unpaired t-test. (C) Expression of *Ifn-α* and *Ifn-β* genes evaluated by RT-qPCR in GA muscle from ACTA1-MCM and MCM;FLEXD mice at 2-6-15 d.p.i.. Values expressed as fold change relative to ACTA1-MCM for each time point. n=4-5. Statistical significance assessed by two-way ANOVA, with Sidak's multiple comparisons test, \*\*p<0.01.

Fig. S5.

Supplementary 5

A

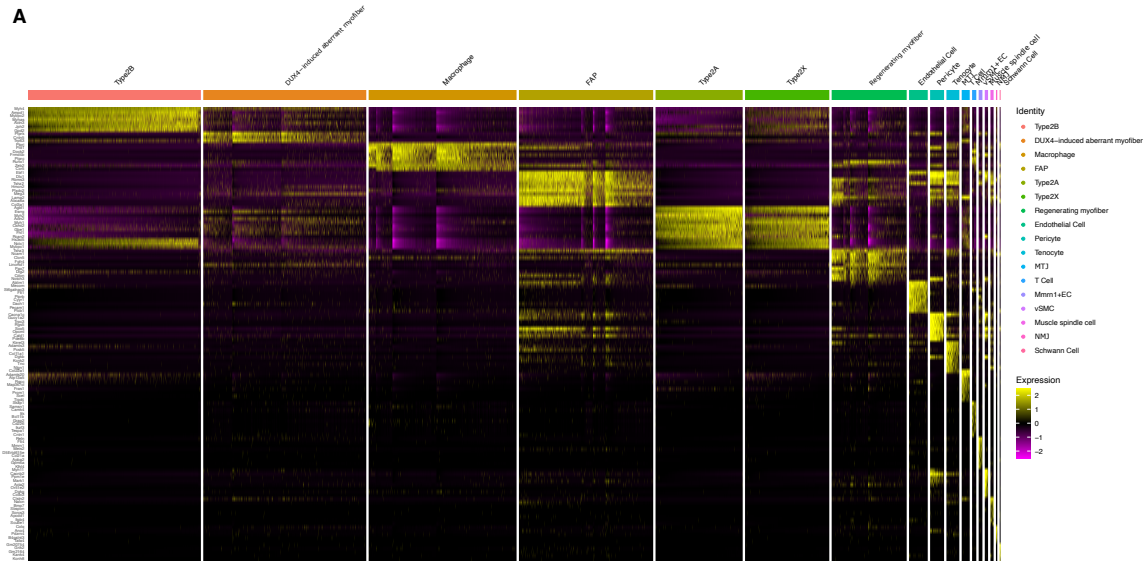

B

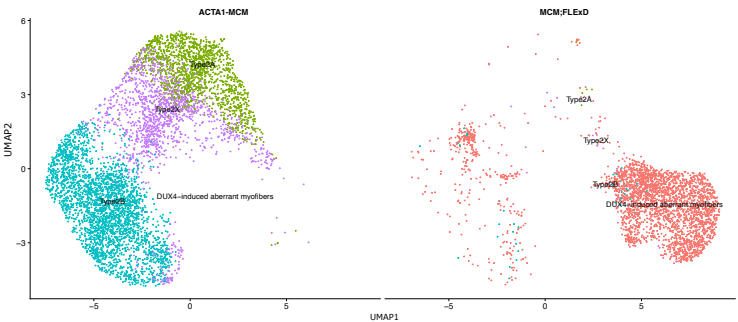

C

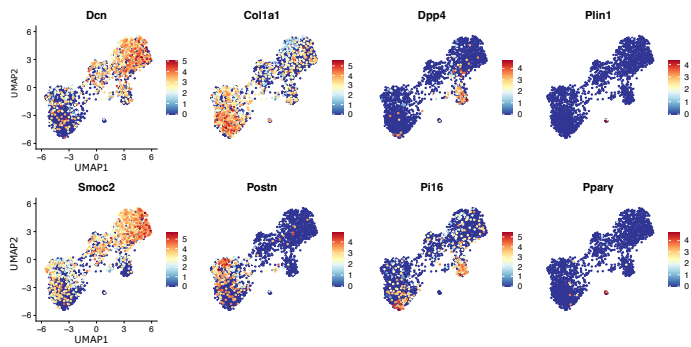

D

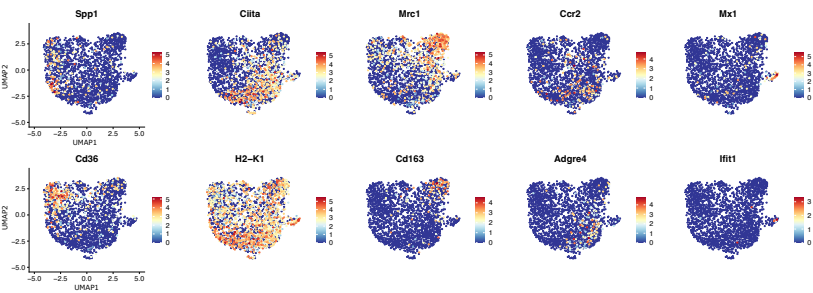

**Figure S5. snRNA-seq cluster-specific gene signatures in ACTA1-MCM and MCM;FLExD muscles**

(A) Heatmap for the top enriched genes per cluster in ACTA1-MCM and MCM;FLExD. (B) UMAP embedding of snRNA-seq data colored by myofiber cluster in ACTA1-MCM and MCM;FLExD. (C) Feature plots showing normalized expression of FAP sub-cluster enriched genes in ACTA1-MCM and MCM;FLExD. (D) Feature plots showing normalized expression of macrophage sub-cluster enriched genes in ACTA1-MCM and MCM;FLExD.

**Fig. S6.**

**Supplementary 6**

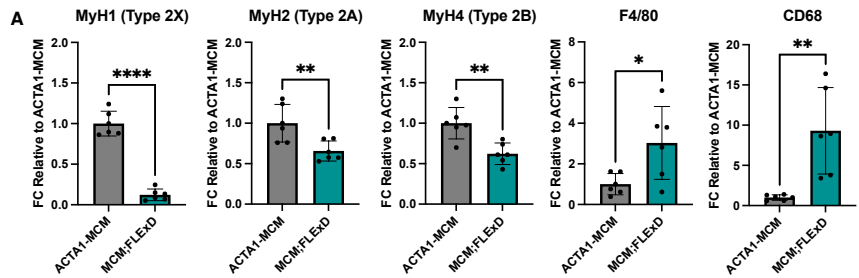

**Figure S6. Validation of the main snRNA-seq results**

(A) RT-qPCR analysis of the expression of Myh1 (marker of Type 2X myofibers), Myh2 (marker of Type 2A myofibers), Myh4 (marker of Type 2B myofibers), F4/80 and CD68 (markers of macrophages) in TA from ACTA1-MCM and MCM;FLEXD mice at 15 d.p.i.. Values expressed as fold change relative to ACTA1-MCM mice. n=6. Statistical significance assessed by Unpaired t-test, \*p<0.05; \*\*p<0.01; \*\*\*\*p<0.0001.

**Fig. S7.**

**Supplementary 7**

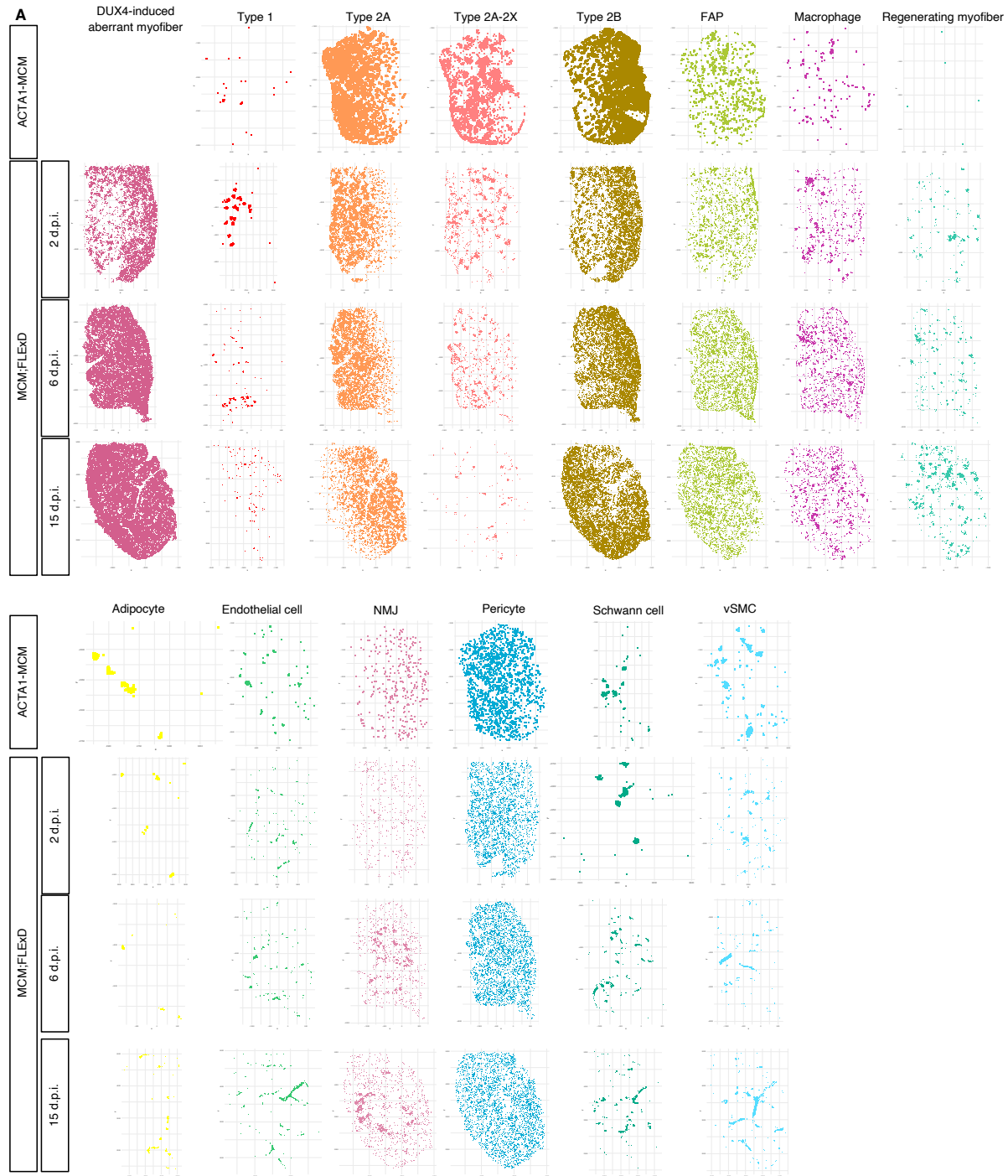

**Figure S7. Spatial location of cell types identified by ST**

(A) Spatial location of the indicated cell types using Visium HD spatial transcriptomics in ACTA1-MCM and MCM;FLEXD muscles at 2, 6, and 15 d.p.i..

Fig. S8.

Supplementary 8

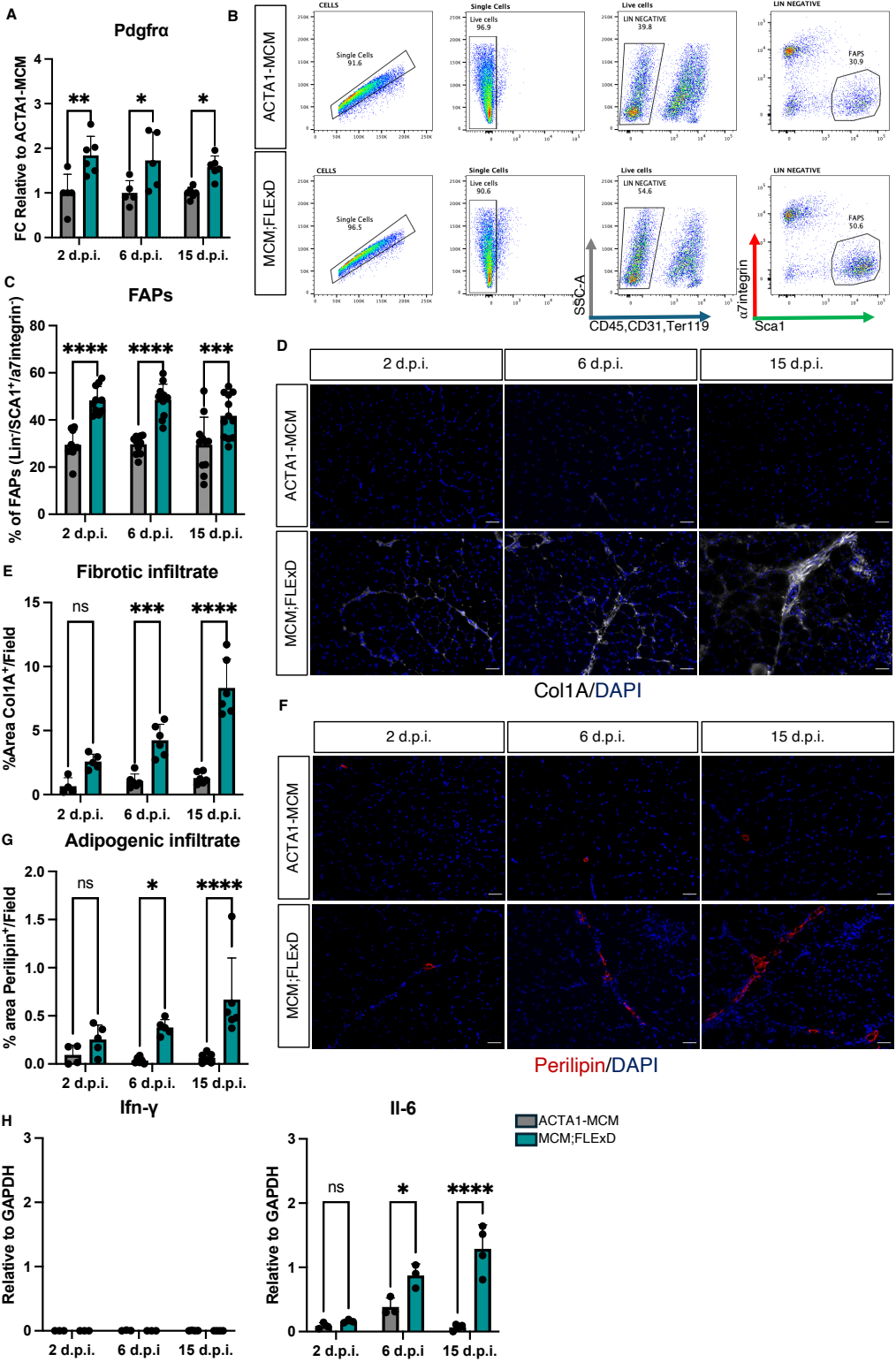

##### Figure S8. Expansion of FAPs in MCM;FLExD mice

(A) *Pdgfra* expression evaluated by RT-qPCR in ACTA1-MCM and MCM;FLExD mice at 2-6-15 d.p.i.. Values expressed as fold change relative to ACTA1-MCM mice for each time point. n=4-6. Statistical significance assessed by two-way ANOVA, with Sidak's multiple comparisons test, \*$p < 0.05$; \*\*$p < 0.01$. (B) Representative FACS sorting strategy for isolating FAPs from ACTA1-MCM and MCM;FLExD mice. FAPs are defined as Lineage negative (Ter119<sup>-</sup>/CD45<sup>-</sup>/CD31<sup>-</sup>) and $\alpha 7$-integrin<sup>-</sup>/Sca1<sup>+</sup> cells. (C) FACS quantification of FAPs from ACTA1-MCM and MCM;FLExD mice at 2-6-15 d.p.i.. n=9-12. Statistical significance assessed by Two-way analysis of variance (ANOVA) with Sidak's multiple comparisons test, \*\*\*$p < 0.001$, \*\*\*\*$p < 0.0001$. (D) Representative images of immunofluorescence for Col1A (grey) and nuclei in DAPI (blue) on TA from ACTA1-MCM and MCM;FLExD mice at 2-6-15 d.p.i.. n=4-6. (E) Quantification of the immunofluorescence shown in D. Statistical significance assessed by Two-way analysis of variance (ANOVA), with Sidak's multiple comparisons test, \*$p < 0.05$, \*\*\*$p < 0.001$, \*\*\*\*$p < 0.0001$. (F) Representative images of immunofluorescence for Perilipin (red) and nuclei in DAPI (blue) on TA from ACTA1-MCM and MCM;FLExD mice at 2-6-15 d.p.i.. n=4-6. (G) Quantification of the immunofluorescence shown in F. Statistical significance assessed by Two-way analysis of variance (ANOVA), with Sidak's multiple comparisons test, \*$p < 0.05$, \*\*\*\*$p < 0.0001$. (H) Expression of the *Ifn-γ* and *Il-6* evaluated by RT-qPCR in FAPs isolated from ACTA1-MCM and MCM;FLExD mice at 2-6-15 d.p.i.. Values expressed as relative to *Gapdh*. n=3-4. Statistical significance assessed by Two-way analysis of variance (ANOVA), with Sidak's multiple comparisons test, \*$p < 0.05$, \*\*\*\*$p < 0.0001$.

**Fig. S9.**

**Supplementary 9**

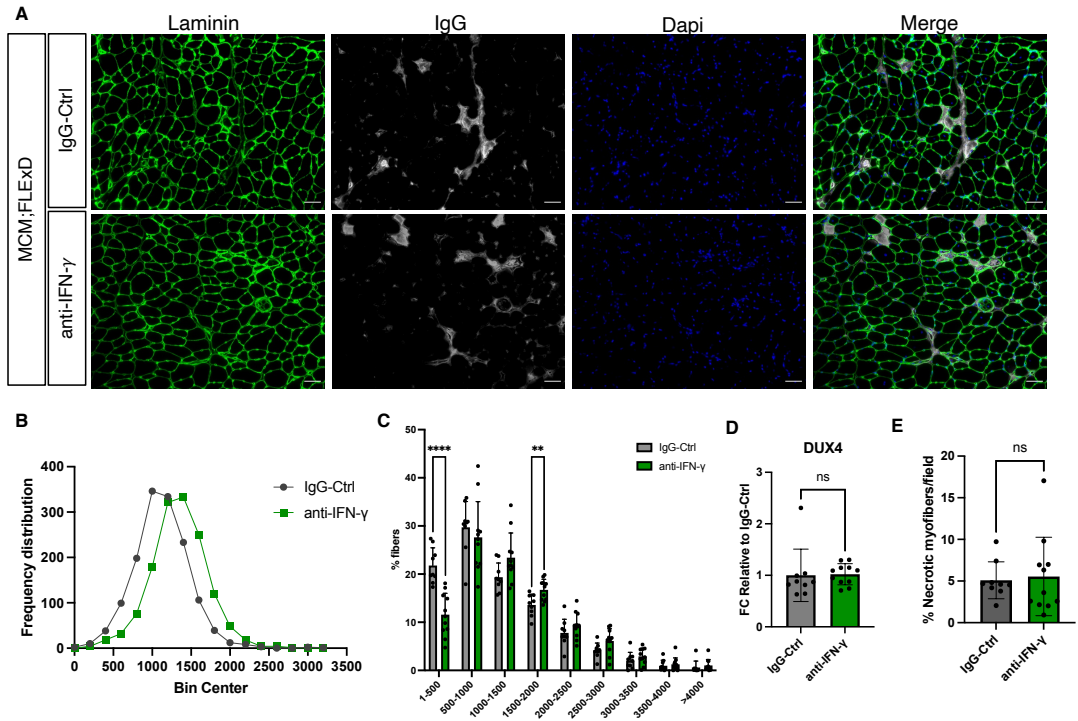

**Figure S9. IFN-γ neutralization preserve muscle size of MCM;FLEXD mice**

(A) Representative images of immunofluorescence for laminin (green), IgG (to detect necrotic fibers, grey) and nuclei in DAPI (blue) on TA from MCM;FLEXD mice treated with IgG-Ctrl or anti-IFN-γ neutralizing antibodies. Scale bar: 50μm. (B) Frequency distribution of the CSA quantification shown in Fig. 7K. (C) Percent of myofibers with the indicated size range of TA from MCM;FLEXD mice treated with IgG-Ctrl (n=9) or anti-IFN-γ (n=11). Statistical significance assessed by Two-way analysis of variance (ANOVA), with Sidak's multiple comparisons test, \*\*p<0.01; \*\*\*\*p<0.0001. (D) RT-qPCR showing the expression of *DUX4* in GA from MCM;FLEXD mice treated with IgG-Ctrl (n=9) or anti-IFN-γ (n=11). (E) Quantification of necrotic myofibers in A. For the graphs in D and E the statistical significance has been assessed by Unpaired t-test.
